# Generation length of the world’s amphibians and reptiles

**DOI:** 10.1101/2024.05.23.595540

**Authors:** G. Mancini, L. Santini, V. Cazalis, F. Ficetola, S. Meiri, U. Roll, S. Silvestri, D. Pincheira-Donoso, M. Di Marco

## Abstract

Variation in life histories influences demographic processes from adaptive changes to population declines leading to extinction. Among life history traits, generation length offers a critical feature to forecast species’ demographic trajectories such as population declines (widely used by the IUCN Red List of Threatened Species) and adaptability to environmental change over time. Therefore, estimates of generation length are crucial to monitor demographic stability or future change in highly threatened organisms, particularly ectothermic tetrapods (amphibians and reptiles) – which rank among the most threatened groups – but for which uncertainty in future impacts remains high. Despite its importance, generation length for amphibians and reptiles is largely missing. Here, we aimed to fill-in this gap by modeling generation lengths for amphibians, squamates and testudines as a function of species size, climate, life history, and phylogeny using generalized additive models and phylogenetic generalized least squares. We obtained estimates of generation lengths for 4,543 (52%) amphibians, 8,464 (72%) squamates and 118 (32%) testudines. Our models performed well for most families, for example Bufonidae in amphibians, Lacertidae and Colubridae in squamates and Geoemydidae in testudines, while we found high uncertainty around the prediction of a few families, notably Chamaeleonidae. Species’ body size and mean temperature were the main predictors of generation length in all groups. Although our estimates are not meant to substitute robust and validated measurements from field studies or natural history museums, they can help reduce existing biases in conservation assessments until field data will be comprehensively available.

## Introduction

Life history variation underlies the whole spectrum of demographic processes, from the pace of adaptive change to the risk of population declines that lead to extinction (Pincheira-Donoso et al., 2021; Ripple et al., 2017). The aggregated influence of life history traits on such demographic processes is often represented by species generation lengths – the average age of parents of the current cohort (i.e., newborn individuals in the population; Charlesworth, 1994; Gingerich, 2019; IUCN Standards and Petitions Committee, 2022). Generation length has been incorporated into the assessment of species conservation status developed by the IUCN Red List of Threatened Species (hereafter IUCN Red List), the most authoritative tool for assessing species extinction risk (Betts et al., 2020). The IUCN Red List classifies extinction risk based on five quantitative criteria, which rely on small ranges, small population size or population decline (IUCN, 2012).

Generation length is used to standardize the time frame over which to measure species decline or probability of extinction (Criteria A, C and E; e.g., decline measured in 3 generations or 10 years, whichever is the longest, for criterion A), ensuring comparability across very different taxa (Mace et al., 2008). Such information is particularly needed to assess extinction risk due to ongoing, or future, environmental changes (Foden et al., 2019; Mancini et al., 2023). Unfortunately, data on generation length are currently missing from most published Red List assessments (76% of all threatened taxa on the Red List; IUCN, 2023), and this can limit assessors ability to accurately apply criteria A, C and E. Indeed, a time frame of 10 years (i.e. the minimum interval allowed for criterion A; IUCN, 2023) is often used in the absence of generation time information.

Several methods to calculate generation length for individual species exist. Life tables are recognized as the most accurate approach (https://www.iucnredlist.org/resources/generation-length-calculator), however the data required to compute these (population-specific information on survival and fecundity) are often missing and difficult to obtain. Other suggested methods to approximate generation length require data on species’ reproductive traits, which are available for many species (e.g. de Magalhaes & Costa, 2009). For example, the lifespan method calculates generation length using age at first reproduction and reproductive lifespan (summing age at first reproduction to the product of a constant *z* with the reproductive lifespan) (IUCN Standards and Petitions Committee, 2022). This method has been used to estimate generation length for all mammals (Pacifici et al., 2013) and birds (Bird et al., 2020), and the results have been widely used in Red List assessments.

Only 2% of amphibians and 4% of reptiles have available generation length data (IUCN, 2023). This major lacuna inevitably influenced the applicability of IUCN Red List criteria for these groups. In fact, only 10% of amphibians and 15% of reptiles have been assessed based on criteria A and C, compared to 53% of mammals and 70% of birds. It is possible that more herptiles could trigger these criteria if generation length was known, particularly if a significant number of species had a long (>3.3 years) generation time. Furthermore, in reptiles, these criteria were mostly used to assess testudines and crocodiles, rather than the more widespread group of squamata (Meiri et al., 2023). Indeed, of 1,978 squamate species assessed as threatened (lizards and snakes) only 169 were based, at least in part, on criteria A or C. We suspect assessors could probably not have assessed many other species based on these criteria, because of the lack of generation length data. That said, among the 534 threatened herptiles assessed by criteria A and C, generation length has been published just for 290 species (IUCN, 2023). Despite the generation length of many small vertebrates might be lower than 3 years and therefore the 10 years time frame can be used to apply criterion A, recent studies suggested that some small herptiles can have generation times much longer than previously thought (Lunghi, 2022).

Herptiles are facing major declines due to highly dynamic threats (Cox et al., 2022; Luedtke et al., 2023). For example, 61% and 28% of amphibians and reptiles on the IUCN Red List, with population trend data available, show population declines (IUCN, 2023), mostly driven by habitat loss and disease for amphibians (Luedtke et al., 2023) and by habitat loss and overexploitation for reptiles (Cox et al., 2022). Additionally, climate change is already a critical threat for amphibians and reptiles and is expected to amplify its impact in the future (Murali et al., 2023; Sinervo et al., 2010; Tingley et al., 2019). The proportion of amphibians and reptiles facing high climate exposure is high, despite the limited use of climate risk in threat assessments (especially for reptiles; Mancini et al., 2023; Meiri et al., 2023). Therefore, estimates of generation length for herptiles are strongly needed.

Here, we provided generation lengths for 4,543 amphibian, 8,464 squamate, and 118 testudine species based on global-scale datasets of their life history traits. We used two modeling techniques: Generalized Additive Model (GAM) and Phylogenetic Generalized Least Square (PGLS). Our results can be key for conservation assessments, inform future species distribution shifts (Mancini et al., 2023), and highlight ecological factors that shape life histories (de Magalhaes & Costa, 2009).

## Methods

We collected generation length data for amphibian, squamate, and testudine species from the IUCN data repository and we complemented these data by calculating generation length using the lifespan method for several additional species. We used morphological, climate, life history, and phylogeny data to model herptiles generation length using two algorithms. We validated our models through different validation approaches and provided models’ performances for each group. Finally, we combined the predictions of the two models. We provided generation length for 4,543 amphibians (all predicted), 8,464 squamates (630 from the lifespan method and 7,834 predicted), and 118 testudines (53 from the lifespan method and 65 predicted).

### Species selection

We considered all the amphibians, squamates, and testudines on the IUCN Red List (7,981 amphibians, 9,933 squamates, and 263 testudines). We excluded crocodiles because most (83% Crocodylia species) already have data on generation length. We also excluded Rhynchocephalia, with only one living representative (the tuatara). As this study can be very important for Red List assessors, we used the IUCN Red List taxonomy as a reference to which we matched the taxonomies of the other databases used in this study. We excluded all amphibians and reptiles species classified as extinct on the IUCN Red List (37 amphibians, 24 squamates and 8 testudines).

### Calculation of generation length from reproductive data

We collected generation length data from the IUCN data repository (IUCN, 2023), which provided information on generation length for 116 amphibians, 150 squamates, and 130 testudines. We complemented these data calculating generation length using the reproductive lifespan method (IUCN Standards and Petitions Committee, 2022):

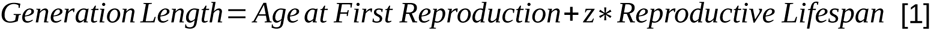

where reproductive lifespan is the difference between maximum longevity and age at first reproduction and *z* is a coefficient representing survivorship and fecundity between young and old individuals. This method has already been used to estimate generation length of the world’s birds (Bird et al., 2020) and mammals (Pacifici et al., 2013) and found to perform best at estimating generation length among the suggested methods on the Red List guidelines (Fung & Waples, 2017). Therefore, we retrieved *z* from [1] as:

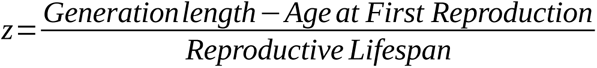

for all species with available data on generation length, age at first reproduction and reproductive lifespan (38 squamates and 39 testudines). We then used the averaged z values, for testudines and squamates separately, to apply [1] on the remaining species with available information only on reproductive lifespan (e.g. the *z* value of squamates was used to calculate generation length of the remaining squamates species through [1]). We could not calculate *z* and apply [1] on amphibians because only 12 species had all three life-history parameters available. Following this step we had generation length data for 116 amphibians (all from IUCN), 880 squamates (150 from IUCN and 630 calculated) and 183 for testudines (130 from IUCN and 53 calculated). Data on age at maturity and maximum longevity were selected from different databases (see below).

### Predictors of generation length

#### Species biological traits selection

We selected biological traits that are expected to be related with generation length and available for most species of herptiles. We used body mass for amphibians and reptiles as we expect generation length to increase with body size (Cooke et al., 2018; Healy et al., 2019). The most recorded data on body size in amphibians is snout-vent length (SVL), however it does not allow comparisons across amphibian taxa (Santini et al., 2018). Thus, for amphibians with unavailable data on body mass, we transformed SVL in body mass using allometric formulas from Santini et al. (2018). These formulas are available only for anurans and salamanders, therefore we excluded all Gymnophiona with no data on body mass from our analysis (n = 17). For amphibians we also used life-history modes from Liedtke et al. (2022) which represent broad categories of reproductive modes and life cycles, such as aquatic, semi-terrestrial, terrestrial, direct development, live-bearing, and paedomorphism. We expect these to influence generation length. For example, paedomorphism is an adaptation to inhospitable environments and associated with long lifespan (e.g. *Proteus*). For squamates we also selected information on insularity, as island reptiles have been found to live generally longer than reptiles on the mainland (Stark et al., 2018). Traits were retrieved from different databases: SVL, body mass, age at maturity and maximum longevity for amphibians from Lucas et al. (2023), Etard et al. (2020), and from unpublished data of the Global Amphibian Biodiversity Project (GABiP); body mass, age at maturity, maximum longevity and insularity for squamates and testudines from Etard et al. (2020) and Meiri (2024).

#### Climate predictors selection

We selected primary climatic variables (mean precipitation, minimum temperature, maximum temperature) from CHELSA database (CHELSA V2.1; Karger et al., 2017, 2021) for each month from the year 1989 to 2018 (resolution: 30 arc seconds). We converted all climate rasters into a Mollweide equal area projection to ensure area remains constant at different latitudes during spatial calculations. We used those primary climate variables to calculate 4 bioclimatic variables: mean annual temperature (bio01), temperature seasonality (bio04), as temperature influences reptiles and amphibians reproductive ecology and metabolism (Shine, 2005; Taylor et al., 2021), mean annual precipitation (bio12) and precipitation seasonality (bio15). Those variables explain most of the climatic variation at global scale (Buckley & Jetz, 2008; Raz et al., 2024) and other factors such as summer and winter temperatures exhibit significant correlations with linear combinations of these 4 variables. We retrieved species’ distribution from the IUCN Red List spatial data repository, which provided distribution information for 5,657 amphibians, 9,203 squamates and 121 testudines. For reptiles with unavailable data in the IUCN Red List spatial data repository, we used polygons from the Global Assessment of Reptiles Distribution (de Oliveira Caetano et al., 2022; Roll et al., 2017), adding 270 range polygons (144 squamates and 126 testudines). We converted the species’ range vector polygons into rasters of ∼1 km at the resolution, matching the resolution of the bioclimatic rasters. We then extracted the 30-year average of the 4 bioclimatic variables from each species’ range.

#### Phylogeny

We used phylogenetic eigenvectors to account for latent traits and phylogenetic relatedness (Diniz-Filho et al., 1998). Phylogenetic trees are comprehensively available only for amphibians and squamates, hence we did not include eigenvectors for testudines. We obtained 100 phylogenetic eigenvectors from the phylogenetic tree for amphibians in Jetz & Pyron (2018) and for squamates in Tonini et al. (2016).

### Statistical Modeling of generation length

We excluded all species with missing data for biological traits (either morphology or life history) or distribution. Our final dataset included 4,680 amphibians, 8,614 squamates and 251 testudines. We modeled generation length, separately for each group, using Generalized Additive Models (GAMs) and Phylogenetic Generalized Least Squares (PGLS). We used the same variables for GAMs and PGLS. We log_10_-transformed generation length, body mass, and precipitation data, and scaled all predictors before running the model. In both models we considered non linear responses between predictors and the response variable. We excluded all collinear predictors with Variance Inflation Factors (VIF) > 3. We selected the best GAM model using the null space penalization approach (Marra & Wood, 2011) and the best PGLS testing all possible combinations of predictors through dredging and selecting the models based on the Akaike information criterion.

For GAMs we also selected a set of eigenvectors to include in the model. We selected the set of eigenvector that provided the autocorrelation of the residuals <10% (Diniz-Filho et al., 2012). This threshold represents a compromise between the need to account for phylogenetic information in the models and include a limited number of variables in the model to avoid overfitting. We selected 4 phylogenetic eigenvectors for amphibians and 6 phylogenetic eigenvectors for squamates based on this procedure (Appendix S1).

We validated the models performing a leave-one-out cross validation for all groups and a family block validation for reptiles (Roberts et al., 2017). When modeling biological data, it is advisable not to use a random cross validation to ensure independence between testing and training set (Roberts et al., 2017). However, in the small sample of amphibians used in the validation some families represented a high proportion of the training set, creating an imbalance between training and testing and potentially making the family block validation inadequate. For both validations, we calculated the Root Mean Square Error (RMSE) and a Normalized Root Mean Square Error (NRMSE) dividing the average RMSE of the family by the average observed generation length of the family. We then used a NRMSE of 0.5 as a benchmark, indicating an error level equivalent to half of the average generation length of the family and representing predictions useful for applications, compared to flat assumptions (e.g. Red List assessment assuming generation length <3 years). To ease the visualization of the results, we presented the errors relative to the leave-one-out cross validation averaged per family in the main text.

We excluded from the prediction all species that exceeded the range of values for any of the predictors used to fit the model to avoid problems associated with extrapolation beyond the training conditions (Ludwig et al., 2023). Differences in availability of predictors led to a different number of species for which we could predict generation length in GAM and PGLS. For example, in GAM prediction we excluded 414 squamates for which phylogenetic data were unavailable, while in PGLS we predicted generation lengths for these 414 species as the phylogeny was only used to calculate the model coefficients.

Finally, we predicted generation length for 4,300 amphibians, 7,609 squamates, and 65 testudines based on GAM and 4,543 amphibians and 7,834 squamates based on PGLS. We calculated the consensus predictions, estimated as the average of the predicted generation length by GAM and PGLS; if the prediction was available only for one model we used it as the final prediction (for detailed statistical modeling see Appendix S1).

## Results

### Calculation of generation length from reproductive data

The estimate of *z* was 0.438+0.002 (mean+SE) for squamates and 0.460+0.056 for testudines. We calculated generation length, using equation [1] and these z values, for a further 630 squamate species and 53 testudines. Generation length for amphibians (all from IUCN) ranged from 2 to 21 years (mean=5.2, SD=3.9). While generation lengths from IUCN database and calculated through lifespan method for squamates ranged from 0.52 to 30 years (mean=6.3, SD=4.5) but mostly ranged from 1 year to 15 years (5th and 95th quantiles); the sample of generation length for testudines ranged from 7.46 to 100 (mean=25.95, SD=14.22).

### Statistical Modeling of generation length

Our best GAM for amphibians included body mass, annual mean temperature, and two phylogenetic eigenvectors (adjusted R^2^ = 0.482; Table S1); while the PGLS included body mass, temperature seasonality and life history modes (adjusted R^2^ = 0.166; Table S1). The final GAM for squamates included body mass, annual mean temperature, temperature seasonality, precipitation seasonality, insularity and four phylogenetic eigenvectors, and explained the variability of generation length well (adjusted R^2^ = 0.545; Table S2). The PGLS included body mass, annual mean temperature, temperature seasonality, insularity and annual mean precipitation (adjusted R^2^ = 0.217; Table S2). For testudines the GAM included body mass and annual mean precipitation (adjusted R^2^ = 0.277; Table S3).

Variable importance differed among models and taxa (Fig. 1). For amphibians, the contribution of each variable was different between GAM and PGLS. Phylogeny contributed the most to the R^2^ in GAM (62%) followed by mean annual temperature (29%), while the most important variable in PGLS was life history mode (74%, not included in the final GAM) followed by body mass (15%). In squamates, the most important variable was body mass in both GAM and PGLS (respectively 61% and 91% contribution to the R^2^). Phylogeny was among the most important variables in GAM, with a cumulative contribution of 30% to the R^2^. Mean annual temperature was the most important climatic variable in both GAM and PGLS (4.5% and 5.5% respectively), followed by temperature seasonality (3% and 1% respectively). In testudines, body mass was the most important variable with 93% contribution to the R^2^.

**Fig. 1.**
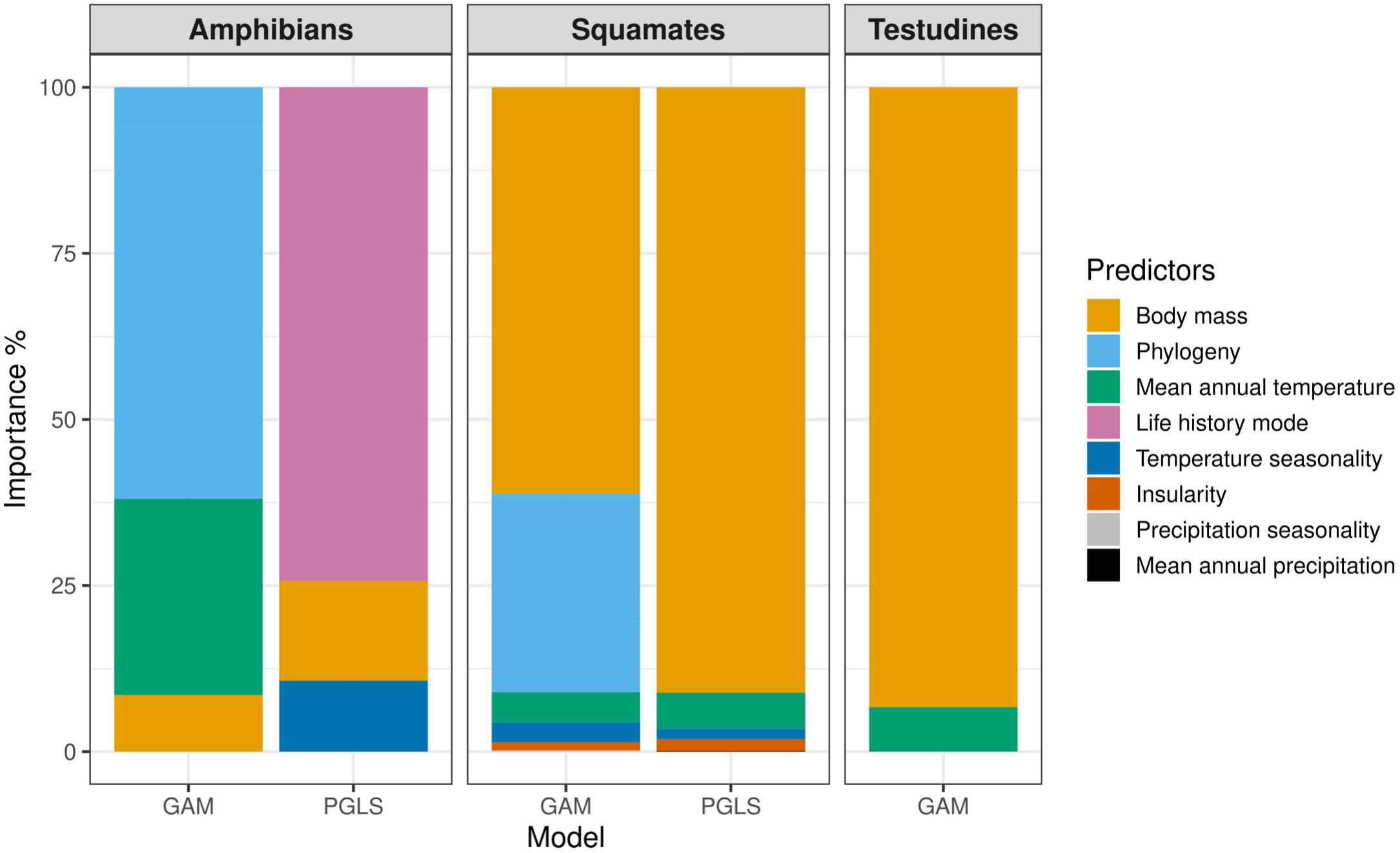
Variable importance. Plot showing the variable importance for GAM and PGLS. Variable importance calculated as the contribution of each variable to the R^2^ of the model. Phylogeny represents the cumulated importance of the phylogenetic eigenvectors (in GAM for amphibians and squamates only). In PGLS phylogeny is accounted for in estimating the model error, thus it had no influence on the R^2^ of the model.

For amphibians, body mass was positively related to generation length in both models (Fig. 2; Fig. S1a,b). In GAM we found annual mean temperature showing a negative relationship with generation length (Fig. 2; Fig. S1b). While, in PGLS we found temperature seasonality showing a positive relationship with generation length (Fig. 2; Fig. S1d) and terrestrial and paedomorphic species showing the highest generation length among the life history modes (Fig. 2; Fig. S1e). For squamates, the responses of GAM and PGLS were overall similar. Body mass had a positive relationship with generation length (Fig. 2; Fig. S2a,f), whereas annual mean temperature showed a negative relationship (Fig. 2; Fig. S2b,g). Temperature seasonality showed a non-linear trend (Fig. 2; Fig. S2c,h) with the lowest generation length for species under high temperature seasonality. Insular species presented higher generation length on average compared to mainland species while marine species showed the highest value of generation length compared to insular and terrestrial species (Fig. 2; Fig. S2d,i). In GAM, precipitation seasonality showed a marginal positive relationship with generation length (Fig. 2; Fig S2e), while in PGLS annual mean precipitation showed a weak negative relationship with generation length of squamates (Fig. 2; Fig. S2j). For testudines, generation length increased with body mass (Fig. 2; Fig. S3a) and decreased with annual mean temperature (Fig. 2; Fig. S3b). Overall, body mass and annual mean temperature were respectively positively and negatively related to generation length across models and taxa. However, the response of generation length varied concerning seasonal climates between amphibians and reptiles.

**Fig. 2.**
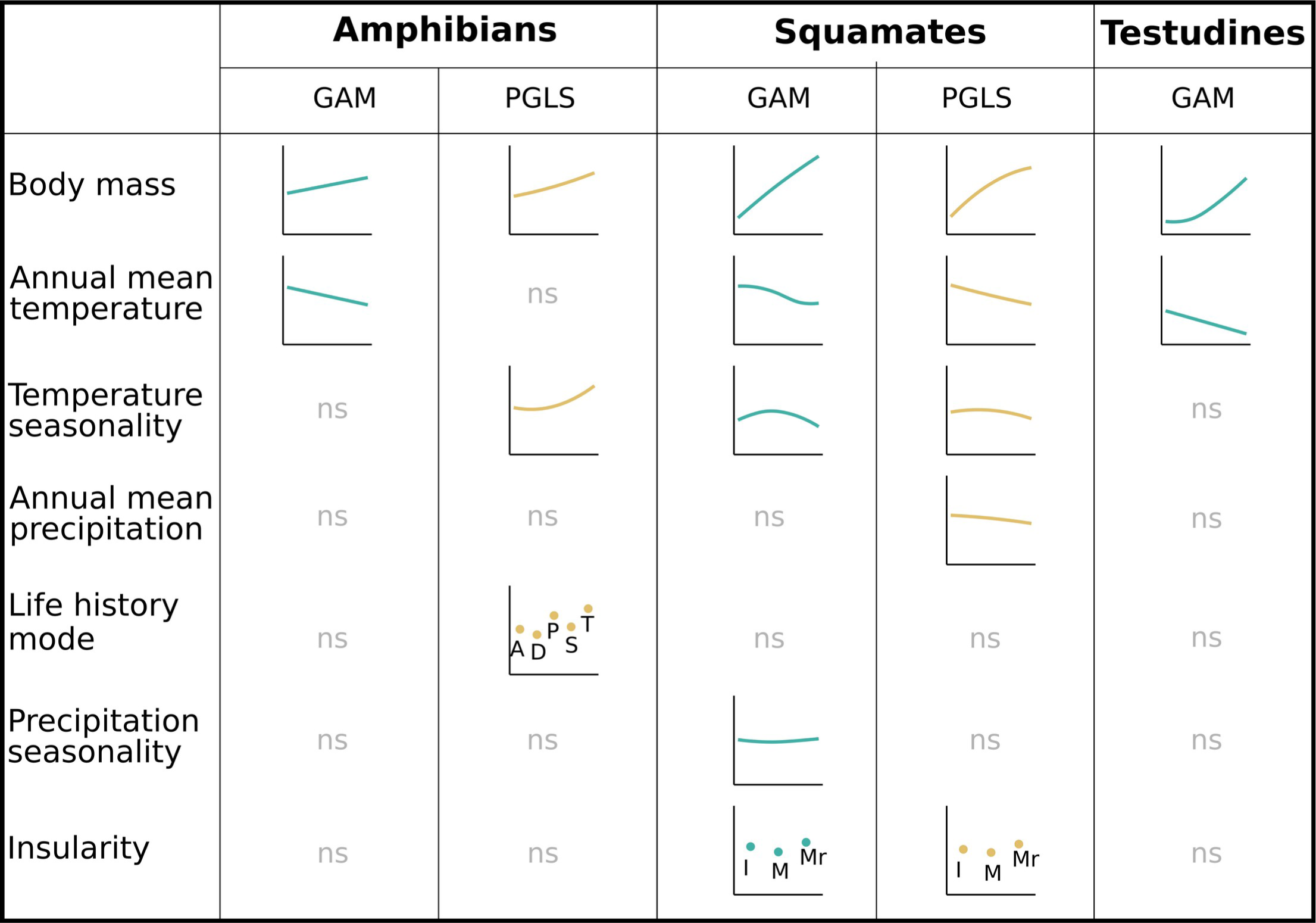
Summary of the relationship between generation length and models’ predictors. Lines represent the predictors response; ns: variable not selected; I: insular; M: mainland; Mr: marine; A: aquatic, D: direct development, P: paedomorphic, S: semi terrestrial, T: terrestrial. For complete responses plots see Fig. S1, Fig. S2 and Fig. S3.

### Validation of GAM and PGLS

For amphibians the NRMSE of the GAM’s leave-one-out validation ranged from 0.07 (Dendrobatidae) to 0.62 (Rhinodermatidae) in GAM and from 0.17 (Ambystomatidae) to 1.45 (Aromobatidae) in PGLS (Fig. 3), with 85% of the families exhibited a NRMSE of 0.5 or lower in GAM and 61% in PGLS. While the NRMSE for squamates ranged from 0.105 (Leptotyphiopidae) to 0.946 (Acrochordidae) for GAM (Fig. 3) and from 0.027 (Lamprophiidae) to 1.07 (Chamaleonidae), with 79% of the families (38 out of 48) exhibited a NRMSE of 0.5 or lower in both GAM and PGLS. Finally, almost all testudines showed a NRMSE <0.5 (Fig. 3), only Dermochelydae showed a NRMSE >0.5 (but the family has just one species).

**Fig. 3.**
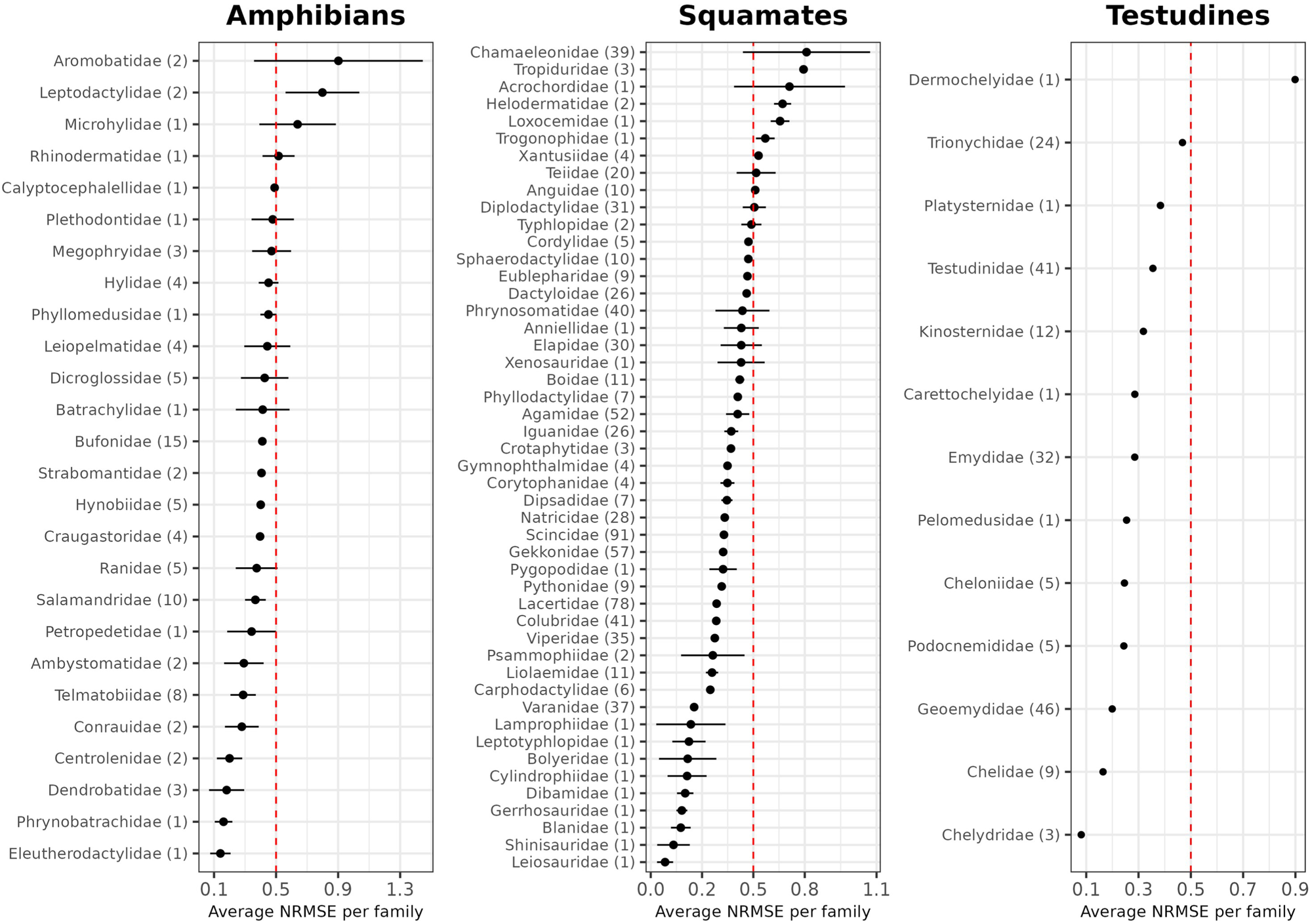
Plot showing the Normalized Root Mean Square Error averaged per family of the leave-one-out validation for GAM (all groups) and PGLS (squamates and amphibians only). Range intervals represent difference bewteen the NRMSE of GAM and PGLS, while points represent the average value. Numbers in brackets next to family names represent the number of species per family. Dashed red line at 0.5 represents the error equal to half of the generation length of the family.

Most amphibia and squamate families showed little variability between validation of GAM and PGLS, while few showed high error uncertainty, notably Aromobatidae for amphibians and Chamaleonidae for squamates (for detailed validation results see Appendix S2).

### Prediction

The generation length predicted by GAM and PGLS for amphibians were not very strongly correlated (Pearson r = 0.43; Fig. S9). The average generation length ranged from 2.68 years to 12.72 years (mean=3.9 years, SD=0.87). We predicted a generation length >10 years only for 6 species: *Proteus anguinus* (12.72 years), *Amphiuma tridactylum* (12.07 years), *Siren lacertina* (11.47 years), *Amphiuma means* (10.88 years), *Siren intermedia* (10.32 years) and *Cryptobranchus alleganiensis* (10.05 years) (Fig 4; Tab. 1). We predicted generation length < 3.3 years for 912 amphibian species (20% of our sample; i.e., for which the 10-year timeframe will still be used for IUCN criteria A, C, and E) (Fig. S10).

**Fig 4.**
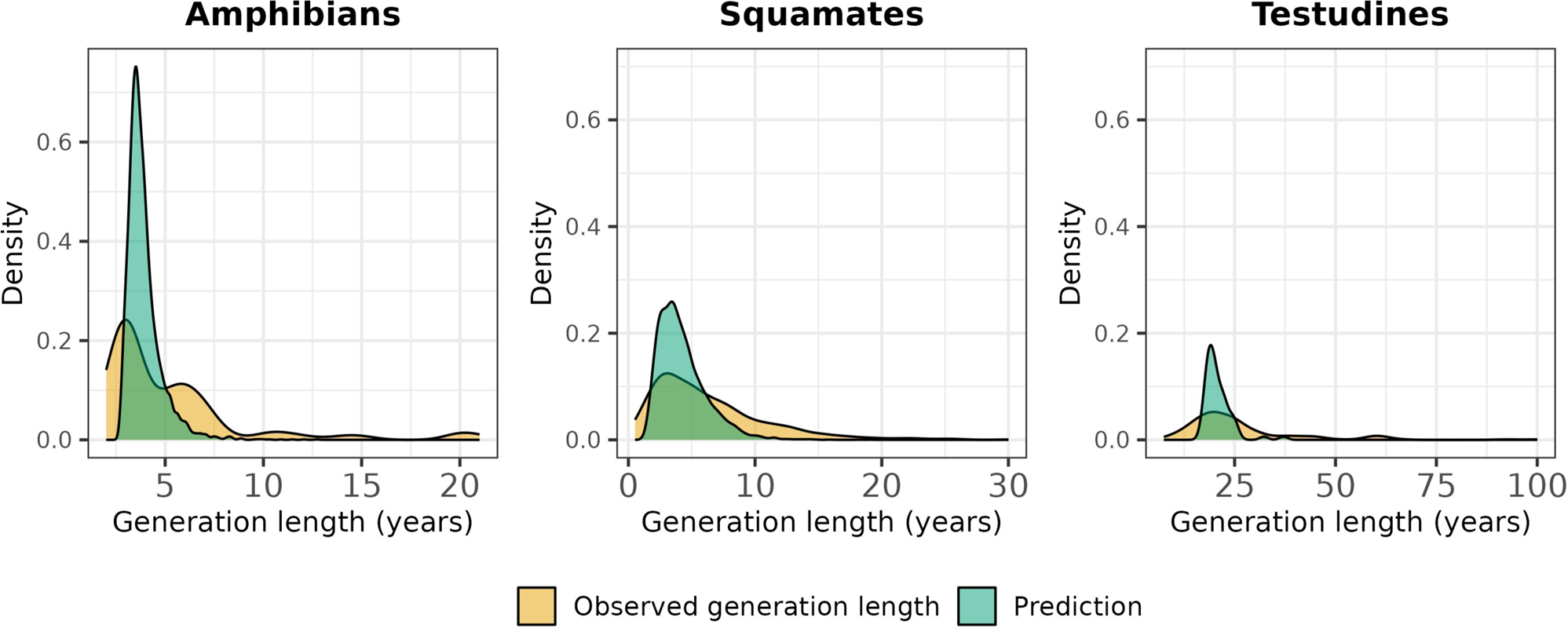
Density plots showing the distribution of the predicted and the observed generation length for 4,659 amphibian, 8,614 squamate and 251 testudines species. Observed generation length included data from IUCN and data calculated using the lifespan method. Predicted generation length refers to the average of GAM and PGLS predictions for squamates and amphibians, and GAM prediction for testudines.

**Table 1.**
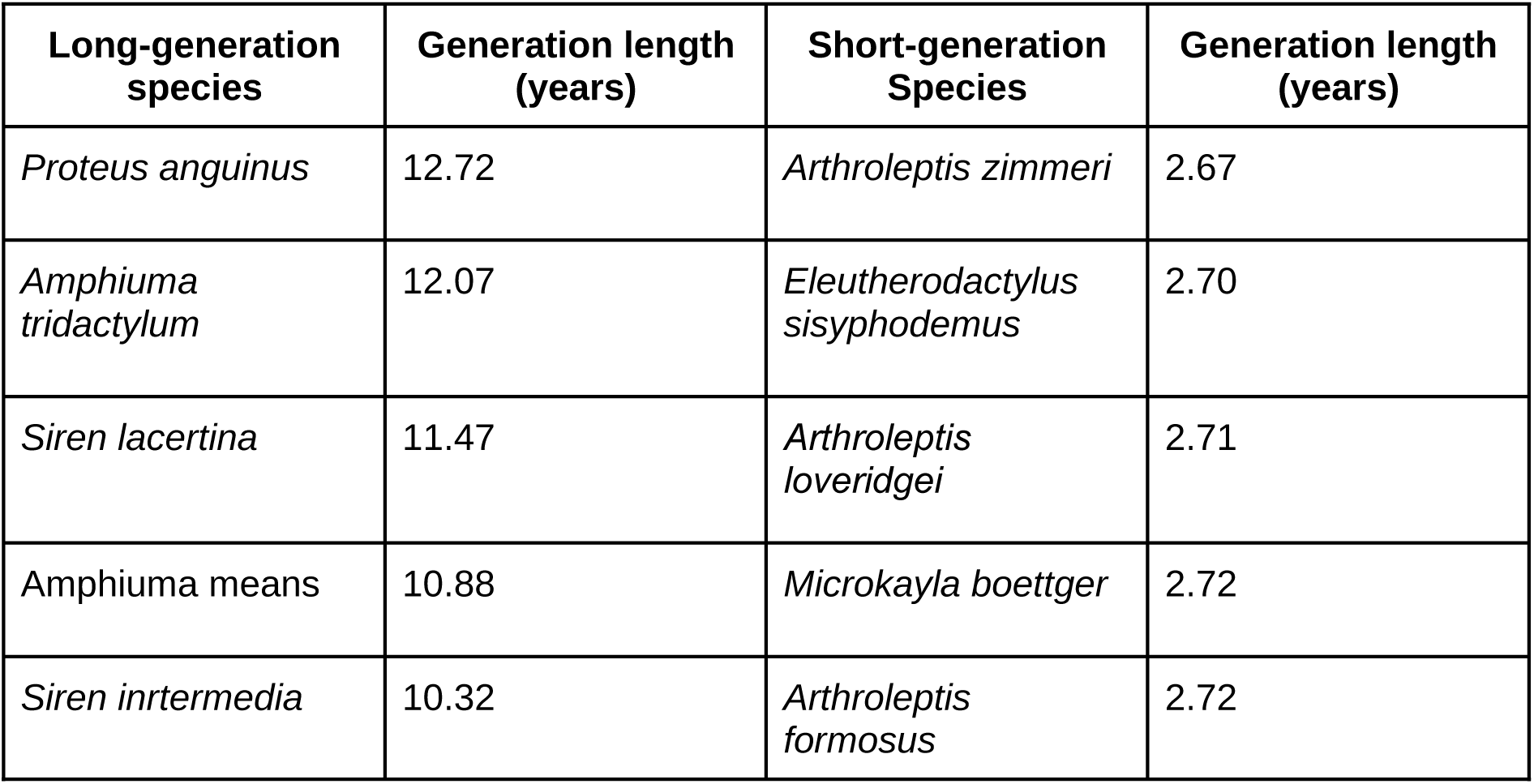
Highest and lowest average predicted generation lengths for amphibians.

The generation lengths predicted by GAM and PGLS for squamates were overall very similar (r = 0.87; Fig. S9). The average predicted generation lengths for squamates ranged from 1.16 to 19.13 years (mean=4.28 years, SD=1.92), with 5,054 (64.5% of our sample) species with generation length >3.3 years (Fig. 4). We predicted the longest generation lengths for families of snakes (Fig. 4; Tab. 2). Notably, 80 of the 104 squamates species for which we predicted a generation length *≥* 10 years were snakes. We predicted a generation length *≤*3.3 years for 2,780 species (35.5%). Among them, 57.2% were represented by four families: Scincidae (23.5%), Gekkonidae (17.9%), Dactyloidae (9.4%), Gymnophthalmidae (6.4%) (Fig. S10): the smallest-bodied of lizards.

**Table 2.** Highest and lowest average predicted generation lengths for squamates.

| **Long-generation species** | **Generation length (years)** | **Short-generation Species** | **Generation length (years)** |
| --- | --- | --- | --- |
| *Hierophis cypriensis* | 19.13 | *Gymnophthalmus cryptus* | 1.15 |
| *Protobothrops himalayanus* | 18.21 | *Amphisbaena slevini* | 1.22 |
| *Philodryas chamissonis* | 15.86 | *Amapasaurus tetradactylus* | 1.25 |
| *Elaphe quatuorlineata* | 15.84 | *Cynisca haugi* | 1.29 |
| *Protobothrops kaulbacki* | 14.89 | *Lepidoblepharis microlepis* | 1.32 |

The predicted generation length for 65 testudines ranged from 17.07 to 37.12 years (mean=20.82 years, SD=3.33) (Fig. S9). We predicted generation length *≥*20 years for 31 species out of 65, 58% represented by Chelidae and Geoemydidae (Fig 4; Tab. 3). While on average the lowest generation length was predicted for Podocnemididae, Pelomedusidae and Kinosternidae (Fig. S10).

**Table 3.** Highest and lowest predicted generation lengths for testudines by GAM.

| **Long-generation species** | **Generation length (years)** | **Short-generation Species** | **Generation length (years)** |
| --- | --- | --- | --- |
| *Chitra indica* | 37.12 | *Kinosternon creaseri* | 17.07 |
| *Elusor macrurus* | 32.25 | *Claudius angustatus* | 17.46 |
| *Amyda cartilaginea* | 25.58 | *Mesoclemmys dahli* | 17.64 |
| *Dermatemys mawii* | 25.27 | *Rhinoclemmys areolata* | 17.65 |
| *Mauremys japonica* | 24.90 | *Malayemys macrocephala* | 17.72 |

## Discussion

In this study we estimated the generation length of 4,543 amphibian, 8,464 squamate, and 118 testudine species, providing new generation length data for 32-72% species in these groups. We found that generation length increased with the body mass of the species and decreased with annual mean temperature in all groups. However, differences among taxa concerned the role of climate seasonality. Despite moderate statistical fit, validation showed that our models performed well for most families across all groups, including some of the most specious families, for example Bufonidae in amphibians, Lacertidae and Colubridae in squamates and Geoemydidae in testudines. At the same time, we found high uncertainty around the prediction of a few families, notably Chamaeleonidae.

We found a consistent positive relationship between body mass and generation length of species across amphibians, squamates, and testudines. Generation length increased with body mass across all groups, which is consistent with previous studies that showed that species of herptiles live generally longer when the size increases (Scharf et al., 2015; Stark et al., 2018; Stark & Meiri, 2018), a pattern similar to the one of birds and mammals (Healy et al., 2014). Similarly, we found a consistent negative effect of annual mean temperature on species generation length across the three taxa. This is again in line with comparative analysis of the influence of climate in herptiles’ life history, showing that herptiles live longer in cold climates than in warm climates (Cabezas-Cartes et al., 2018; Meiri et al., 2013; Stark & Meiri, 2018; Zhang & Lu, 2012). Species living in cold climates exhibit slower metabolism and physiological activities, constrained by limited resources availability and reduced reproductive season (Meiri et al., 2012). Conversely, species living in warm climates tend to reproduce earlier, facilitated by extended reproductive seasons and ample resources availability, which potentially affect the accumulation of harmful metabolic by-products resulting in overall shorter longevity (Scharf et al., 2015; Stark et al., 2018).

On the contrary, the relationship between generation length and climate seasonality differed between amphibians and squamates. Temperature seasonality had a positive effect in amphibians, but a marginal negative effect on generation length of squamates. In climates characterized by a strong seasonality, herptiles are inactive during some parts of the year. Throughout these periods of brumation, the level of activity decreases (Dubiner et al., 2023), prolonging various aspects of species’ life history, such as longevity and generation length (Scharf et al., 2015). We found the highest generation length for terrestrial and paedomorphic species, on average, among the life history modes considered. Paedomorphism is generally associated with inhospitable conditions and scarcity of resources, which can strongly reduce physiological activities and overall result in longer lifespan and generation length (Bonett et al., 2014). The generation lengths of insular endemic, terrestrial and marine squamates were very similar and annual mean precipitation and precipitation seasonality had overall a weak effect on generation length of squamates.

Our results and predictions can have multiple applications for conservation. First, having generation length estimates for amphibians and reptiles can reduce the bias on the application of Red List criteria among tetrapods. Criterion B, which is based on species geographic range and does not require generation length, has been used to classify 80% threatened amphibians and 74% threatened reptiles, compared to only 40% mammals and 20% birds (IUCN, 2023). Although this does not necessarily mean that the species are misclassified, our estimates will enable assessors to take into account different responses to threats for herptiles as done for other vertebrates (e.g. population trends for criteria A and C in addition to changes in geographic range size for criterion B). In fact, our data will be easily accessible for assessors directly in Red List assessment through the sRedList platform (https://sredlist.eu/). Second, our estimates of generation length can improve the accuracy of extinction risk assessments for which a short generation length has been wrongly assumed. Criterion A, for example, measures species responses over a period of 3 generations or 10 years (whichever is the longest). Assessors, based on their expertise, can assume the generation length of a species is lower than 3.3 years and consider 10 years to measure population decline if generation length is not available. This is often assumed for anurans and lizards. However, we found that 73% of anurans and 54% of lizards had a generation length >3.3 years and that their population trends should thus be considered over periods longer than 10 years. For instance, the median of our predicted generation length for Ranidae was 4.6 years and it was 3.46 for Lacertidae (both had good performance in the validation). Furthermore, to apply Criterion C1, other definitions are needed. To grant endangered status under Criterion C1, for example, one must consider a period of 5 years or 2 generations, which will be relevant for species with generations as short as 2.5 years. Moreover, as some threats correlate best with different criteria (e.g., criteria A and C with climate change and land-use change; Henry et al., 2024), assessors can focus on specific species’ responses depending on the sensitivity of the species under study. This might also aid comparative extinction risk modeling for these groups, which has inherent difficulties in predicting new Red List assessments partly due to the limited application of decline-related criteria in the Red List (Di Marco, 2022). Third, our estimates can help the assessment of Data Deficient species on the IUCN Red List, which are particularly numerous in reptiles and amphibians (IUCN, 2023). Generation length can for instance be used to calculate deforestation rates over 3 generations within the species range, that can be sufficient information to reassess a Data Deficient species into a threatened or Near Threatened category (Cazalis et al., 2023; Tracewski et al., 2016). Ultimately, our estimates can also serve as initial parameters for Population Viability Analysis for at-risk species and populations (relevant for criterion E ; Pearson et al., 2014), and in setting goals for on-the-ground conservation actions (beyond the IUCN Red List categories), until true observations of generation lengths will be obtained.

Generation length estimates can also support predictions of the impact of future climate change on herptiles species. Species characterized by long generation length are generally more vulnerable to climate change, as generation length is as a proxy of sensitivity to this threat (Foden et al., 2019; Pearson et al., 2014). Herptiles are ectothermic species that are severely impacted by climate change (Biber et al., 2023; Buckley et al., 2012; Luedtke et al., 2023; Newbold, 2018; Trull et al., 2018), unfortunately, the uncertainty around the potential responses of herptiles to climate change is still high, mostly related to the unavailability of the data needed to measure these responses (Winter et al., 2016). For example, several studies measured species responses to climate over a fixed period instead of a biological meaningful time frame (e.g. 2050 or 2080; Biber et al., 2023; Newbold, 2018). Using such fixed time period may underestimate climate adaptability of many short generation length species (e.g., we found lizards having 3.5 years of generation length on average) and hamper evaluation of future climate change impact with respect to extinction risk assessments (Mancini et al., 2023) or in diversity loss predictions. Similarly, generation length allows to define the number of expected reproductive (dispersal) events within a fixed time frame, which is needed for range shift predictions (Bateman et al., 2013; Santini et al., 2016; Travis et al., 2013). Our estimates can play a crucial role in climate change adaptability studies.

Finally, our estimates can also inform macroecological and evolutionary studies. As life histories generally follow altitudinal or latitudinal gradient (Jetz et al., 2008; Meiri et al., 2020; Morrison & Hero, 2003; Pincheira-Donoso et al., 2021), our estimates can potentially inform studies linking life histories and the geographical patterns, as well as inform studies on the potential influence of life histories on the evolutionary process at global scale (Liedtke et al., 2018).

Our results should be interpreted with caution. First, the data needed to calculate generation length are extremely scarce for herptiles. While morphological data are almost complete, longevity and maturity data are almost completely missing (e.g., 95% and 90% of missing data for longevity for amphibians and reptiles respectively). Although our approach relied on well known patterns, such as allometric and climate relationships (large species and species living in cold climates live longer on average), data as longevity or age at maturity, are crucial to calculate generation length (Bird et al., 2020; IUCN Standards and Petitions Committee, 2022). Our models were conservative in respect of observed data and reluctant to predict both low and high values of generation length, potentially underestimating or overestimating generation length for some species. This may also suggest that observed data are significantly biased with respect to what most species are. These options are not mutually exclusive. In fact, we could only model a small subset of species, due to the amount of missing data. This was particularly problematic for amphibians, for which we only had 87 species to train our models (this only represents ∼1% of the species in this group). Some families were heavily underrepresented (e.g. Eleutherodactylidae or Lamprophiidae). Such families generally exhibited the highest and lowest errors, due to underfitting and overfitting of our models, while well represented families generally performed better. We used the fitted models to predict generation lengths of many species (4,543 amphibians and 7,834 squamates). This might have led to inconsistent predictions, especially for families not included in the validation, although we only included species within the range of values of predictors used in the validation to avoid extrapolation. Therefore, our estimates should be used considering the error but also if the family was included in the validation, and family size. For example, our models performed particularly well for squamates with long generation length. We predicted the longest generations for Iguanidae, Pythonidae, Colubridae, and Viperidae. All these families were above the third quartile of generation length distribution in the input data, had a relatively low NRMSE (<0.5) in both GAM and PGLS, and were well represented in the dataset used in the validation. Conversely, for Chamaeleonidae and Aromobatidae, our models did not perform well (NRMSE>0.8).

All in all, our study should not deter important and much needed additional data collection on the reproductive traits of herptiles, which will allow us to further refine predictions in the future; in the meanwhile predictions for these groups, as produced here, should be favored over simplistic assumptions about generation time values being below 3 years when unknown. Our estimates of generation length can have multiple applications spanning from conservation assessments to evolutionary and biogeographical studies. This new information will help have a comprehensive overview of extinction risk and the ecology of herptiles.

