## Supplementary material for "Generation length of the world’s amphibians and reptiles"

**Appendix S1**

**Statistical Modeling of generation length**

We excluded all species with missing data for predictors or distribution. Our final dataset included 4,659 amphibians, 8,614 squamates and 251 testudines. We modeled generation length, separately for each group, using Generalized Additive Models (GAMs) and Phylogenetic Generalized Least Squares (PGLS). Due to lack of phylogenetic information, we could not run PGLS for testudines. We used GAMs to capture nonlinear responses between predictors and the response variable, setting a gaussian family and using smooth functions and relative smoothing penalty to prevent overfitting (Pedersen et al., 2019). We modeled continuous variables using thin plate splines with k=4 to limit the smooth functions to quadratic and cubic polynomials. To calibrate the penalization on the smooth terms, we set the gamma parameter = 1.4, which has been suggested as a good compromise for reducing overfitting while not excessively penalizing model fit (Boonman et al., 2020; Jiménez-Valverde et al., 2021). We fitted all GAM models using the restricted maximum likelihood (REML). We used PGLS incorporating second order polynomials and we optimized the branch length transformations through maximum likelihood (Freckleton et al., 2002).

We used the same variables for GAMs and PGLS. We $\log_{10}$-transformed generation length, body mass, and precipitation data, and scaled all predictors before running the model. For each group, we tested collinearity among predictors and excluded all those with Variance Inflation Factors (VIF) > 3. Then, we modeled generation length as a function of body mass, climate, and phylogenetic eigenvectors (for amphibians and squamates only and in GAMs, see below), life-history modes for amphibians only, and insularity for squamates only. For GAMs we used the null space penalization approach to select the best model (Marra & Wood, 2011). This approach constructs an extra penalty for each smooth term to remove non-significant variables from the model. We iteratively used this approach to identify non-significant smoothed variables that we successively used as linear predictors in our models, while we removed other non-significant linear predictors. We repeated this process until we found a set of significant variables, either smoothed or linear. While for PGLS we tested all possible combinations of predictors through dredging and selected the best model based on the Akaike information criterion.

For GAMs we selected a set of eigenvectors to include in the models for amphibians and squamates among the 100 extracted. We aimed to select the eigenvectors more strongly related to generation length. Therefore we iteratively added one eigenvector at a time to a GAM model with body mass, climate variables, life-history modes for amphibians only and insularity for squamates only. We then selected the eigenvector that minimized the autocorrelation of the residuals (Diniz-Filho et al., 2012). We repeated this process until we found the set of eigenvectors that provided the autocorrelation of the residuals <10%. This threshold represents a compromise between the need to account for phylogenetic information and include a limited number of variables in the model, as having too many predictor variables with respect to the sample size might lead to overfitting. We selected 4 phylogenetic eigenvectors for amphibians and 6 phylogenetic eigenvectors for squamates based on this procedure (Table S1,2). The PGLS explicitly accounts for the phylogenetic relatedness of species in deriving the coefficients of the model so we did not include the phylogenetic eigenvectors in this model.

We performed a leave-one-out cross validation for all groups and a family block validation for reptiles (Roberts et al., 2017). The samples used in the validation differed between groups due to the respective amount of missing data in the predictors. We could use 87 amphibian (29 species with missing data among the 116 with generation length available), 755 squamate species (125 species with missing data among the 880 with generation length available) and 181 testudines (2 species with missing data among the 183 with generation length available). For the leave-one-out validation we iteratively ran our models on a training set consisting of all species except one, that we used as a testing set. For the family block validation we ran our models on a training set consisting of all families except one that we used as a testing set. When modeling biological data, it is advisable not to use a random cross validation to ensure independence between testing and training set (Roberts et al., 2017). However, the sample we used for amphibians was very small (n = 87) and some families represented a high proportion of the training set, creating an imbalance between testing and training set and potentially making the family block validation inadequate (e.g. Bufonidae represented 17% of the whole sample). We assessed models performance by calculating the Root Mean Square Error (RMSE) and the Normalized Root Mean Square Error (NRMSE). For the leave-one-out validation we calculated the NRMSE dividing the RMSE by the observed value of generation length and a NRMSE averaged per family dividing the average RMSE of the family by the average generation length of the family (i.e. we got a NRMSE per species and a NRMSE per family). While for the family block validation we only calculated the NRMSE per family. We used a NRMSE of 0.5 as a benchmark, indicating an error level equivalent to half of the average generation length of the family and representing predictions useful for applications compared to flat assumptions (e.g. Red List assessment assuming generation length <3 years). To ease the visualization of the results, we presented the errors relative to the leave-one-out cross validation averaged per family in the main text.

We excluded from the prediction all species that exceeded the range of values for any of the predictors used to fit the model to avoid problems associated with extrapolation beyond the training conditions (Ludwig et al., 2023). This led to a different number of species for which we could predict generation length due to the different predictors used in GAM and PGLS. For example, in GAM prediction we excluded 414 squamates for which phylogenetic data were unavailable, while in PGLS we predicted generation lengths for these 414 species as the phylogeny was only used to calculate the model coefficients. Finally, we predicted generation length for 4,300 amphibians, 7,609 squamates, and 65 testudines based on GAM and 4,543 amphibians and 7,834 squamates based on PGLS. We finally calculated the consensus predictions, estimated as the average of the predicted generation length by GAM and PGLS; if the prediction was available only for one model we used it as the final prediction. All predictions were provided with the respective RMSE and NRMSE of both validation methodologies.

We also ran a model for squamates, as described above, including data on viviparity (Meiri, 2024). Viviparity is an adaptation to inhospitable cold climates (Zimin et al., 2022) that may influence generation length. However, we decided to exclude viviparity as the predictions for squamates with and without viviparity were strongly correlated (r=0.98; Fig. S11), it had little influence on generation length (Fig. S12), and we would have had to exclude 1,069 squamates from prediction due to unavailable data on viviparity.

All spatial analysis were performed in GRASS GIS 7.8.6 (GRASS Development Team, 2017) and all statistical analysis were performed in R 4.2.1 (R Core Team, 2023) using RStudio 2023.6.0.421 (Posit Team, 2023) with the following packages: “ape” (Paradis & Schliep, 2019), “caper” (Orme et al., 2023), “ggplot2” (Wickham, 2016), “mgcv” (Wood, 2003, 2004, 2006, 2011; Wood et al., 2016), “MuMin” (Bartoń, 2023), “patchwork” (Pedersen, 2023), “PVR” (Santos, 2018), “tidyverse” (Wickham et al., 2019), “usdm” (Naimi et al., 2014).

**Appendix S2**

**Validation of GAM and PGLS**

For amphibians the RMSE of the GAM's leave-one-out validation ranged from 0.06 to 15.79 years (mean=1.72, SD=2.36; Appendix S11), while for PGLS it ranged from 0.12 to 14.98 (mean=2.45, SD=2.67; Appendix S11). Almost all families of amphibians showed a RMSE <5 years in GAM and PGLS (ranging from a minimum of 0.21 for Dendrobatidae to maximum of 4.94 years for Leiopelatidae in GAM and from 0.62 years for Eleutherodactylidae to 4.54 years for Rhinodermatidae in PGLS). Only Calyptocephallelidae (5.1 years in GAM and 5.22 in PGLS), Rhinodermatidae (6.81 years) in GAM and Leiopelmatidae (9.91 years) in PGLS showed a RMSE > 5 years (Fig. S4). The NRMSE ranged from 0.07 (Dendrobatidae) to 0.62 (Rhinodermatidae) in GAM and from 0.17 (Ambystomatidae) to 1.45 (Aromobatidae) in PGLS (Fig. 3), with 85% of the families exhibited a NRMSE of 0.5 or lower in GAM and 61% in PGLS.

For squamates, the RMSE of the GAM's leave-one-out validation ranged from 0.002 to 21.51 years (mean=2.25, SD=2.49; Appendix S11), while for the PGLS it ranged from 0.0015 to 24.07 years (mean=2.57, SD=2.59; Appendix S11). Overall, the average RMSE of most squamate families was <5 years in both GAM and PGLS (ranging from a minimum of 0.21 for Leptotyphiopidae to maximum of 4.79 years for Iguanidae in GAM and from 0.18 for Lamprophiidae to 4.72 years for Anguidae in PGLS). Four families showed an average RMSE greater than 5 years GAM (i.e. Anguidae, 5.05; Acrochordidae, 5.48;  Loxocemidae, 12.5 and Helodermatidae, 14.5) and four families in PGLS (Diplodactylidae, 5.22; Iguanidae, 5.69; Loxocemidae, 10.8 and Helodermatidae, 12.7) (Fig. S4). The NRMSE of the leave-one-out validation averaged per family ranged from 0.105 (Leptotyphiopidae) to 0.946 (Acrochordidae) for GAM (Fig. 3) and from 0.027 (Lamprophiidae) to 1.07 (Chamaleonidae), with 79% of the families (38 out of 48) exhibited a NRMSE of 0.5 or lower in both GAM and PGLS.

For testudines the RMSE of GAM ranged from 0.001 to 63.43 years (mean=7.97, SD=9.27; Appendix S11). The average RMSE per family ranged from 2.7 years (Chelydridae) to 27 years (Dermochelydae), with 8 families out of 13 showing a RMSE <10 years (Fig. S4). Almost all families of testudines showed a NRMSE <0.5 (Fig. 3), only Dermochelydae showed a NRMSE >0.5 (but the family has just one species).

Overall, we did not find a clear relationship between NRMSE and generation length (Fig. S5) for the three groups neither in GAM or PGLS, in fact short generation length species did not show relatively better performances compared to long generation length species in all groups. However, the absolute error was higher when generation length increased, mostly due to underestimation of generation length for species with long generations (Fig. S6).

The family block validation for reptiles showed higher error on average (Fig. S7,8). The GAM's RMSE for squamates ranged from 0.21 years (Leptotyphlopidae) to 14.73 years (Helodermatidae) (mean=3.54 years; SD=2.86; Fig. S7), while in PLGS it ranged from 0.17 (Lamprophiidae) to 13.06 years (Helodermatidae) (mean=3.14 years; SD=2.63; Fig. S7) in PGLS. The GAM's NRMSE ranged from 0.1 (Leptotyphlopidae) to 1.59 (Phrynosomatidae) and from 0.02 (Lamprophiidae) to 1.02 (Chamaeleonidae) for PGLS, with 48.9% of the families showing a NRMSE of 0.5 or lower for GAM and 57% in PGLS (Fig. S8). The RMSE for testudines ranged from 2.65 years (Chelydridae) to 26.96 years (Dermochelydae) (mean=10.64 years; SD=7.31; Fig. S7). The NRMSE for testudines ranged from 0.08 (Chelydridae) to 0.89 (Dermochelydae) with 77% of families showing a NRMSE <0.5 (Fig. S8).

Table S1. Variable selected in the model for amphibians.

| Variable name | Variable used in final model | |
|---|---|---|
| | GAM | PGLS |
| Body mass (g) | yes | yes |
| Annual mean temperature (K) | yes | no |
| Temperature seasonality | no | yes |
| Annual mean precipitation (mm/month) | no | no |
| Precipitation seasonality | no | no |
| Life history modes | no | yes |
| Eigenvector 8 | no | - |
| Eigenvector 62 | no | |
| Eigenvector 71 | yes | |
| Eigenvector 99 | yes | - |

Table S2. Variable selected in the model for squamates.

| Variable name | Variable used in final model | |
|---|---|---|
| | GAM | PGLS |
| Body mass (g) | yes | yes |
| Annual mean temperature (K) | yes | yes |
| Temperature seasonality | yes | yes |
| Annual mean precipitation (mm/month) | no | yes |
| Precipitation seasonality | yes | no |
| Insularity | yes | yes |
| Eigenvector 3 | yes | - |
| Eigenvector 18 | no | - |
| Eigenvector 22 | yes | - |
| Eigenvector 28 | yes | - |
| Eigenvector 35 | no | - |
| Eigenvector 87 | yes | - |

Table S3. Variable selected in the model for testudines.

| Variable name | Variable used in final model |
|---|---|
| | GAM |
| Body mass (g) | yes |
| Annual mean temperature (K) | yes |
| Temperature seasonality | no |
| Annual mean precipitation (mm/month) | no |
| Precipitation seasonality | no |

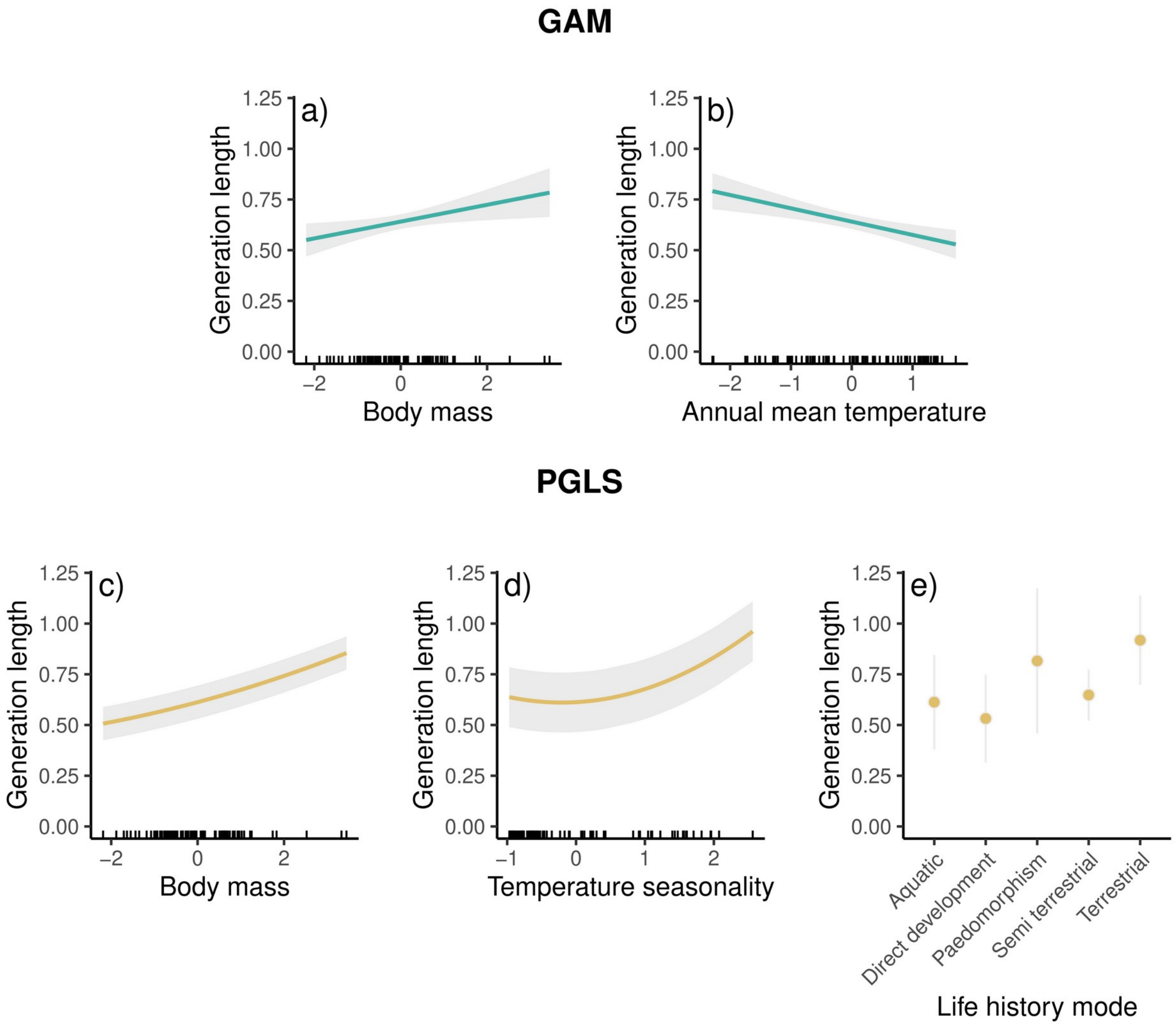


*Fig S1. Amphibians partial dependence plots displaying the relationship between generation length and body mass, climate variables and life history modes. All numerical variables were standardized. Shaded area represented standard error.*

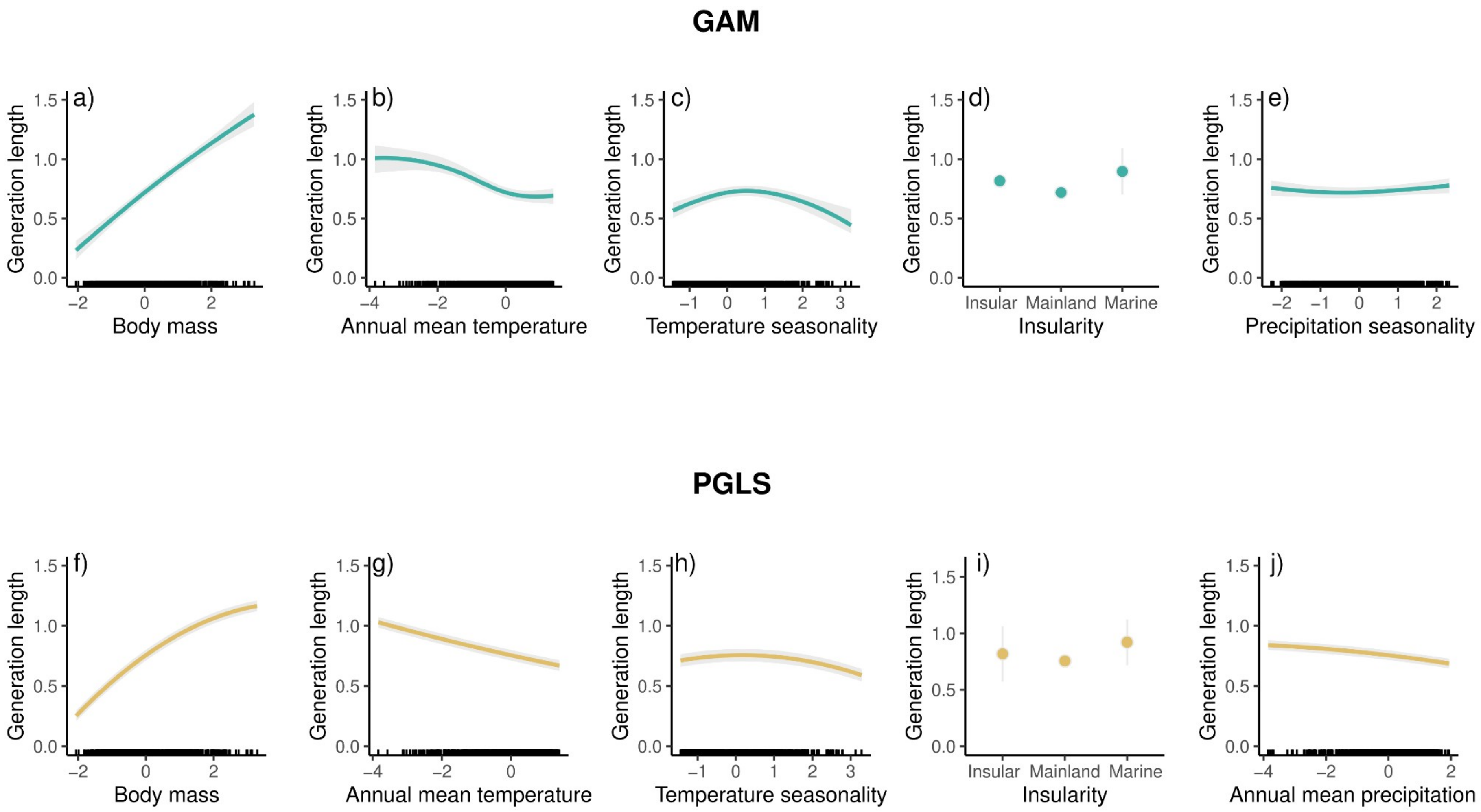


*Fig S2. Squamates partial dependence plots displaying the relationship between generation length and body mass, climate variables and insularity. All numerical variables were standardized. Shaded area represented standard error.*

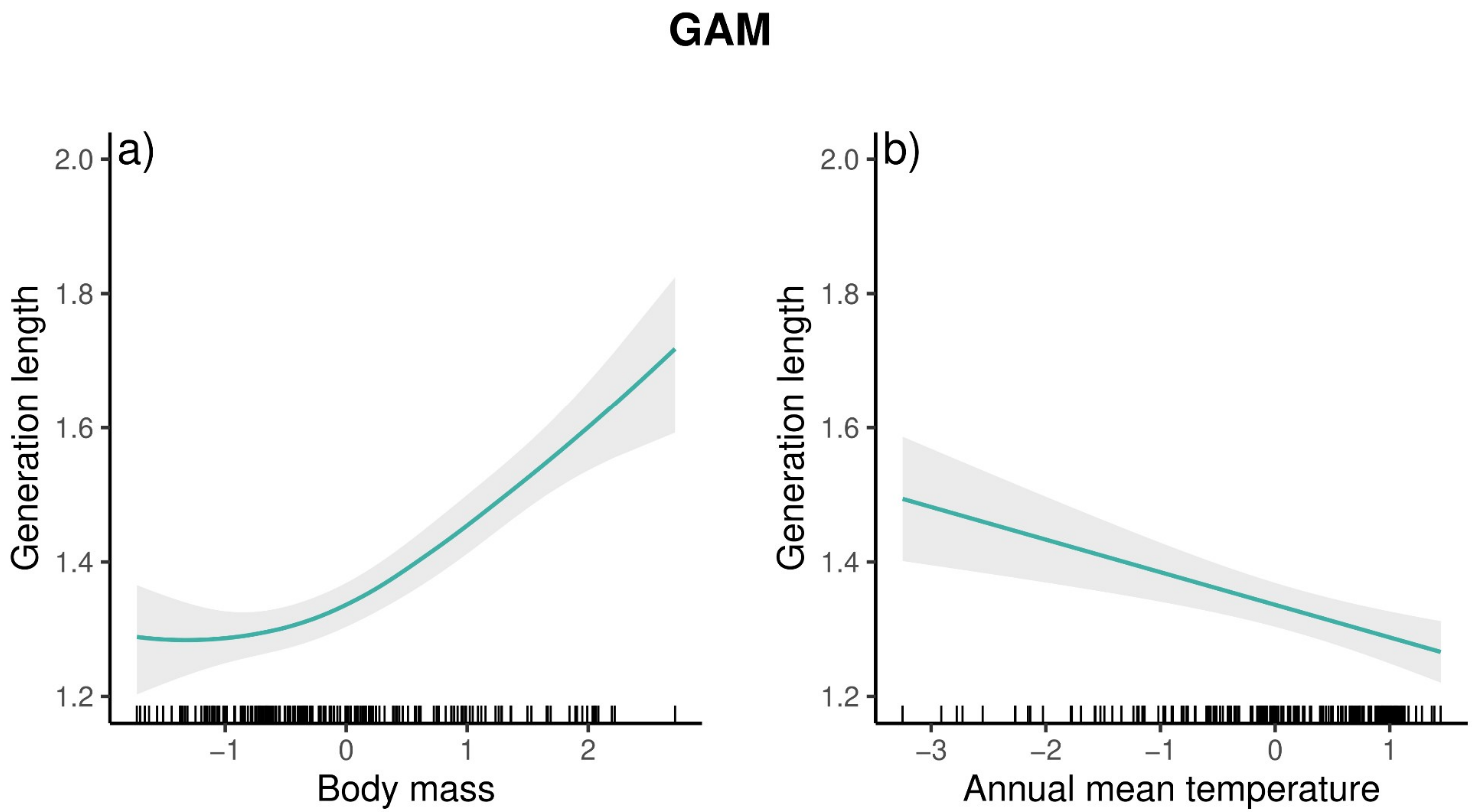


*Fig S3. Testudines partial dependence plots displaying the relationship between generation length and body mass, annual mean temperature. All numerical variables were standardized. Shaded area represented standard error.*

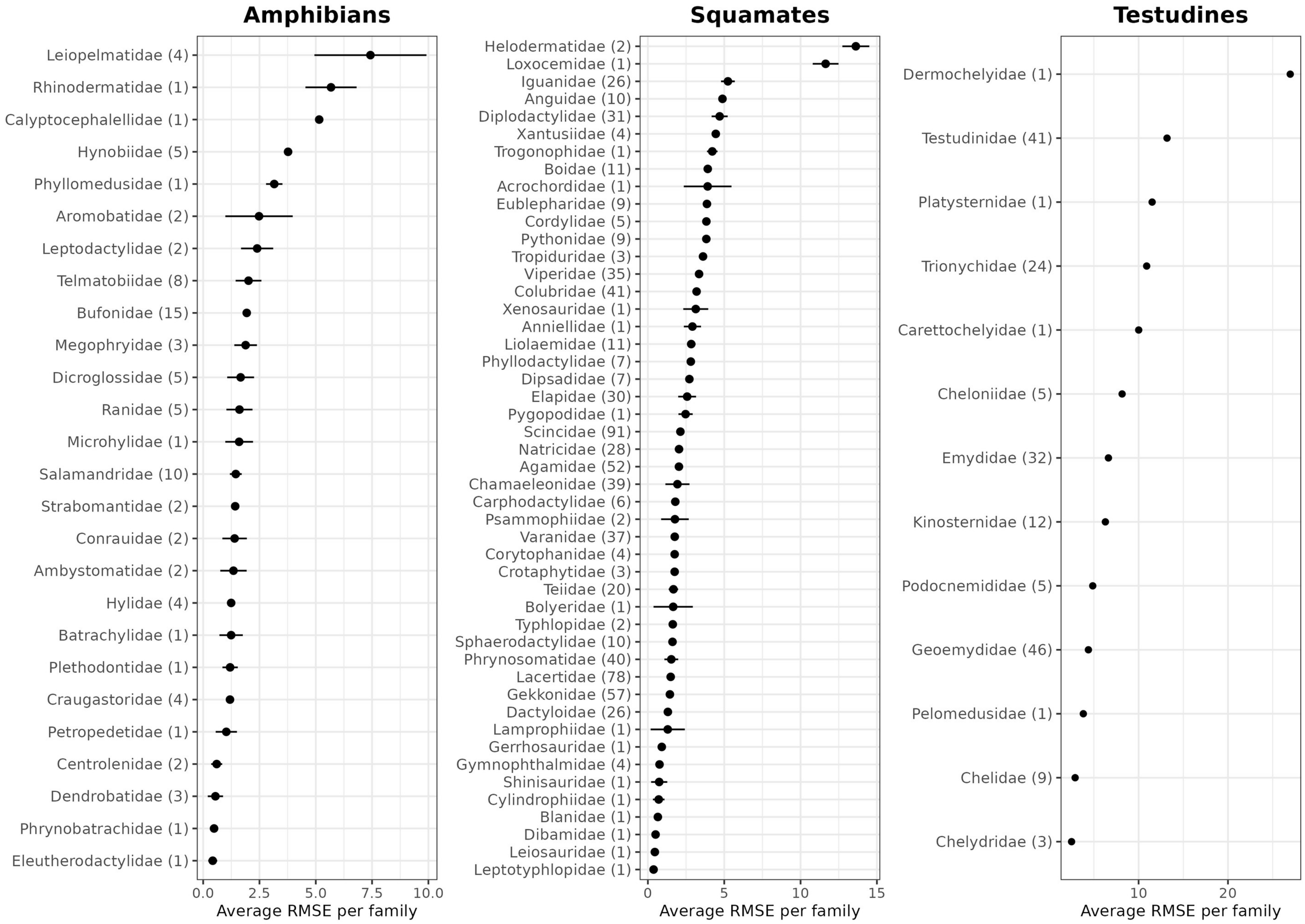
Amphibians
Leiopelmatidae (4)
Rhinodermatidae (1)
Calyptocephalellidae (1)
Hynobiidae (5)
Phyllomedusidae (1)
Aromobatidae (2)
Leptodactylidae (2)
Telmatobiidae (8)
Bufonidae (15)
Megophryidae (3)
Dicroglossidae (5)
Ranidae (5)
Microhylidae (1)
Salamandridae (10)
Strabomantidae (2)
Conrauidae (2)
Ambystomatidae (2)
Hylidae (4)
Batrachylidae (1)
Plethodontidae (1)
Craugastoridae (4)
Petropedetidae (1)
Centrolenidae (2)
Dendrobatidae (3)
Phrynobatrachidae (1)
Eleutherodactylidae (1)
0.0
2.5
5.0
7.5
10.0
Average RMSE per family
Squamates
Helodermatidae (2)
Loxocemidae (1)
Iguanidae (26)
Anguidae (10)
Diplodactylidae (31)
Xantusiidae (4)
Trogonophidae (1)
Boidae (11)
Acrochordidae (1)
Eublepharidae (9)
Cordylidae (5)
Pythonidae (9)
Tropiduridae (3)
Viperidae (35)
Colubridae (41)
Xenosauridae (1)
Anniellidae (1)
Liolaemidae (11)
Phyllodactylidae (7)
Dipsadidae (7)
Elapidae (30)
Pygopodidae (1)
Scincidae (91)
Natricidae (28)
Agamidae (52)
Chamaeleonidae (39)
Carphodactylidae (6)
Psammophiidae (2)
Varanidae (37)
Corytophanidae (4)
Crotaphytidae (3)
Teiidae (20)
Bolyeridae (1)
Typhlopidae (2)
Sphaerodactylidae (10)
Phrynosomatidae (40)
Lacertidae (78)
Gekkonidae (57)
Dactyloidae (26)
Lamprophiidae (1)
Gerrhosauridae (1)
Gymnophthalmidae (4)
Shinisauridae (1)
Cylindrophiidae (1)
Blanidae (1)
Dibamidae (1)
Leiosauridae (1)
Leptotyphlopidae (1)
0
5
10
15
Average RMSE per family
Testudines
Dermochelyidae (1)
Testudinidae (41)
Platysternidae (1)
Trionychidae (24)
Carettochelyidae (1)
Cheloniidae (5)
Emydidae (32)
Kinosternidae (12)
Podocnemididae (5)
Geoemydidae (46)
Pelomedusidae (1)
Chelidae (9)
Chelydridae (3)
10
20
Average RMSE per family

*Fig S4. Plot showing Root Mean Square Error averaged per family of the leave-one-out validation for GAM (all groups) and PGLS (squamates and amphibia only). Range intervals represent the RMSE of GAM and PGLS, while points represent the average value. Numbers near family names represent respective family size.*

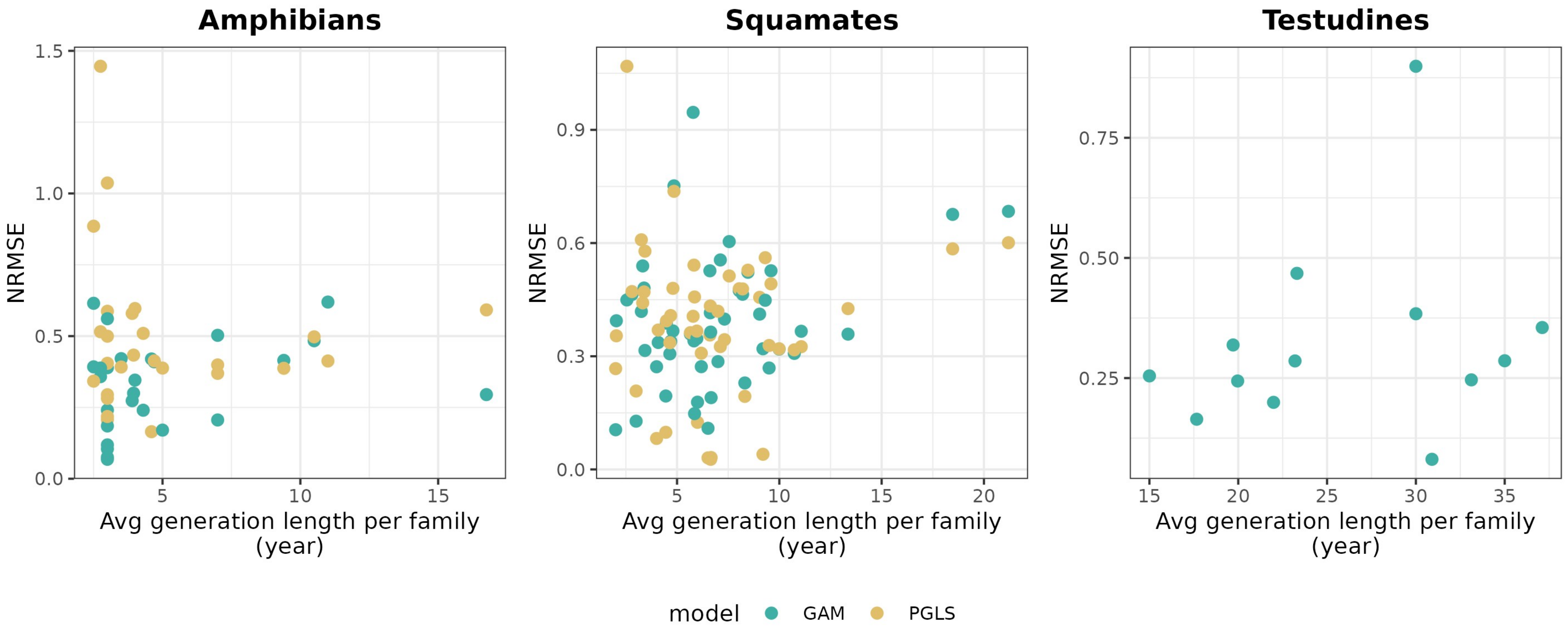


*Fig S5. Plot showing relationship between NRMSE of the leave one out validation (y axis) and generation length (x axis) both averaged per family .*

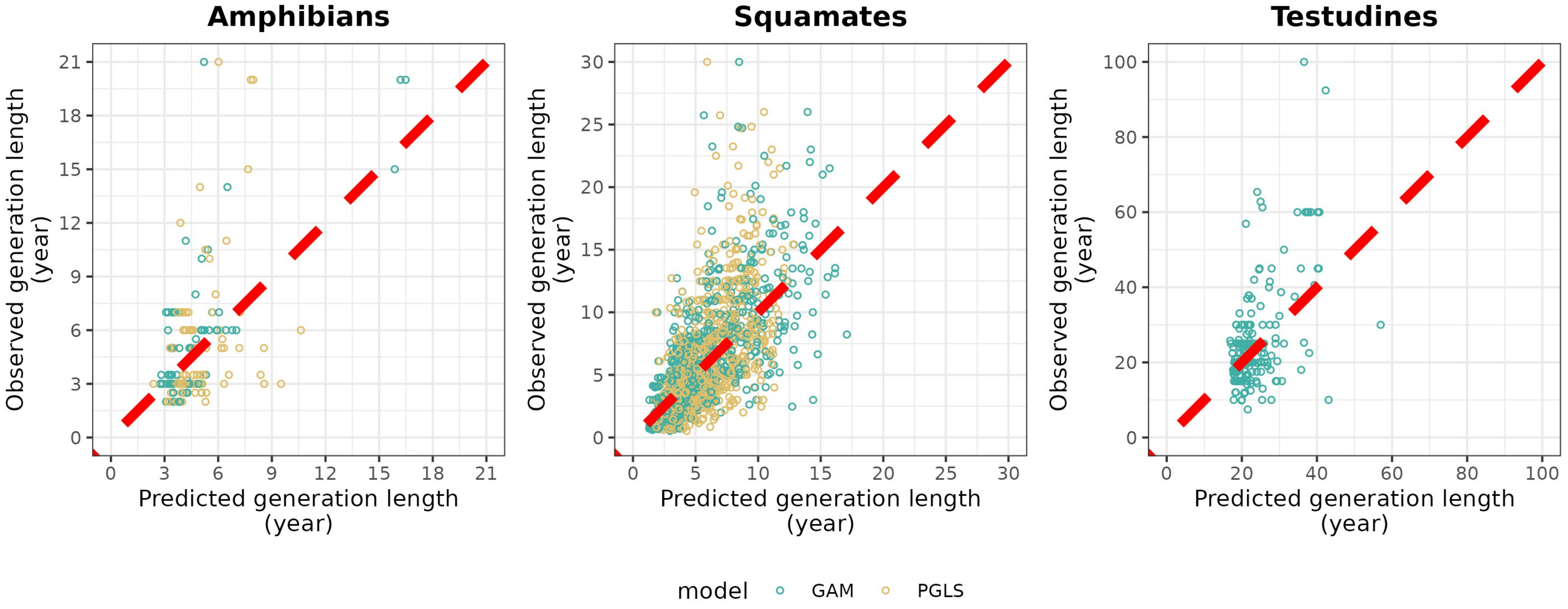


*Fig S6. Scatterplot representing the relationship between observed generation length (y axis) and predicted generation length by GAM or PGLS (x axis). The red dashed line represents the bisector. The points close to the bisector represent little difference between observed and predicted generation length, while points far from the bisector represent a great difference between observed and predicted generation length.*

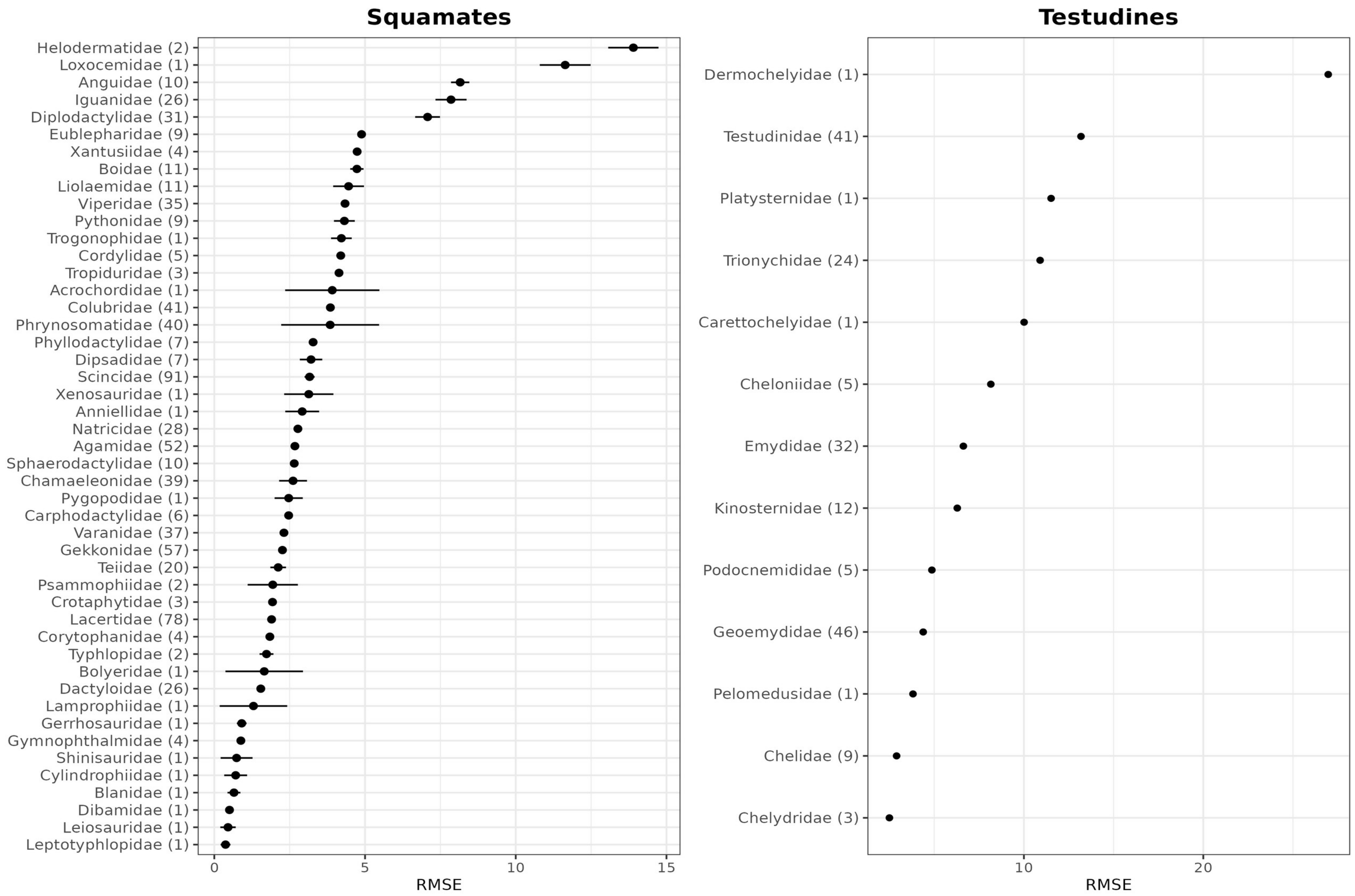

Squamates
Testudines
Helodermatidae (2)
Loxocemidae (1)
Anguidae (10)
Iguanidae (26)
Diplodactylidae (31)
Eublepharidae (9)
Xantusiidae (4)
Boidae (11)
Liolaemidae (11)
Viperidae (35)
Pythonidae (9)
Trogonophidae (1)
Cordylidae (5)
Tropiduridae (3)
Acrochordidae (1)
Colubridae (41)
Phrynosomatidae (40)
Phyllodactylidae (7)
Dipsadidae (7)
Scincidae (91)
Xenosauridae (1)
Anniellidae (1)
Natricidae (28)
Agamidae (52)
Sphaerodactylidae (10)
Chamaeleonidae (39)
Pygopodidae (1)
Carphodactylidae (6)
Varanidae (37)
Gekkonidae (57)
Teiidae (20)
Psammophiidae (2)
Crotaphytidae (3)
Lacertidae (78)
Corytophanidae (4)
Typhlopidae (2)
Bolyeridae (1)
Dactyloidae (26)
Lamprophiidae (1)
Gerrhosauridae (1)
Gymnophthalmidae (4)
Shinisauridae (1)
Cylindrophiidae (1)
Blanidae (1)
Dibamidae (1)
Leiosauridae (1)
Leptotyphlopidae (1)
0
5
10
15
RMSE
Dermochelyidae (1)
Testudinidae (41)
Platysternidae (1)
Trionychidae (24)
Carettochelyidae (1)
Cheloniidae (5)
Emydidae (32)
Kinosternidae (12)
Podocnemididae (5)
Geoemydidae (46)
Pelomedusidae (1)
Chelidae (9)
Chelydridae (3)
10
20
RMSE

*Fig. S7. Plot showing Root Mean Square Error performance averaged of the family block validation for GAM (all groups) and PGLS (squamates and amphibia only). Range intervals represent the RMSE of GAM and PGLS, while points represent the average value. Numbers near family names represent respective family size.*

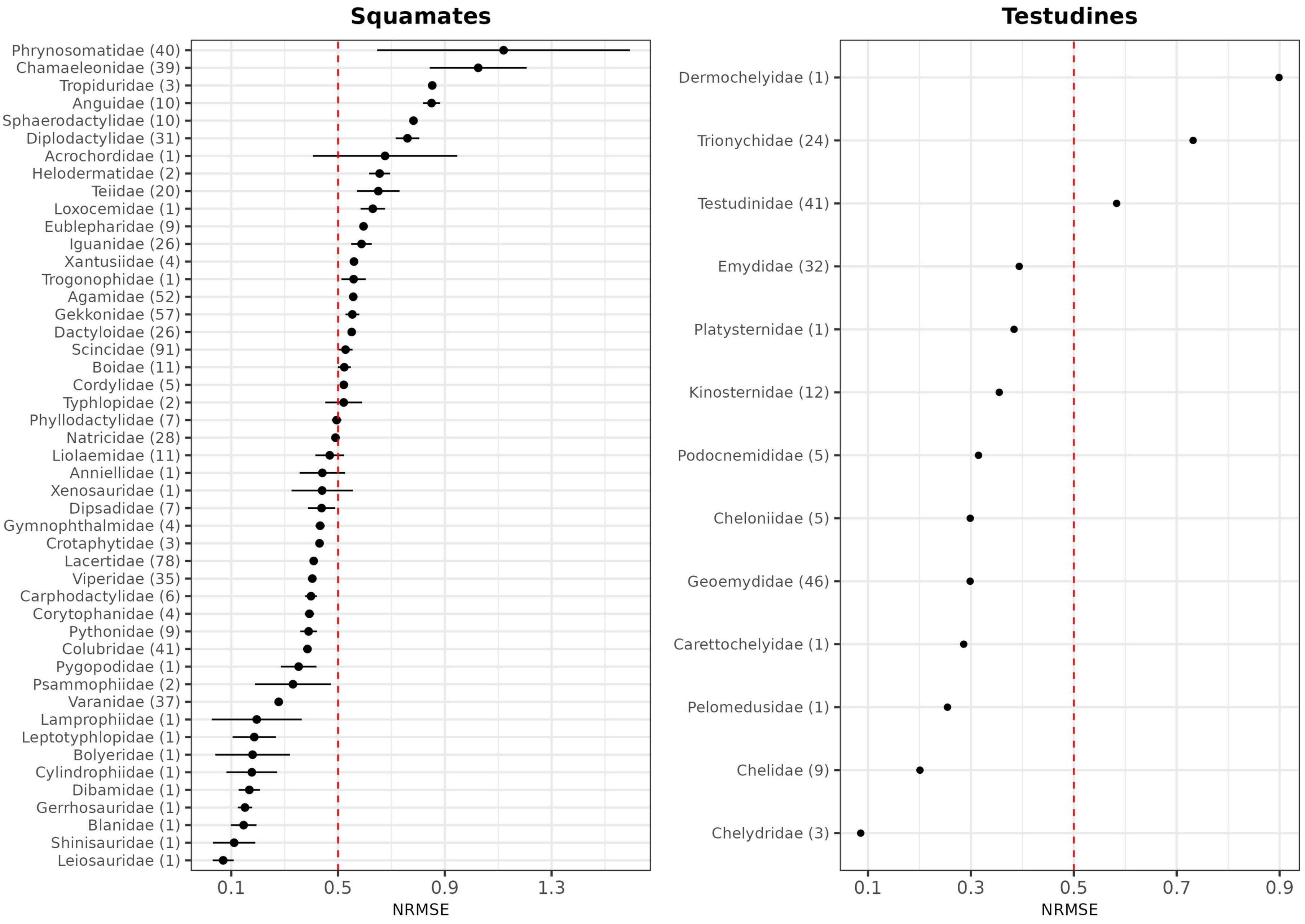
Squamates
Phrynosomatidae (40)
Chamaeleonidae (39)
Tropiduridae (3)
Anguidae (10)
Sphaerodactylidae (10)
Diplodactylidae (31)
Acrochordidae (1)
Helodermatidae (2)
Teiidae (20)
Loxocemidae (1)
Eublepharidae (9)
Iguanidae (26)
Xantusiidae (4)
Trogonophidae (1)
Agamidae (52)
Gekkonidae (57)
Dactyloidae (26)
Scincidae (91)
Boidae (11)
Cordylidae (5)
Typhlopidae (2)
Phyllodactylidae (7)
Natricidae (28)
Liolaemidae (11)
Anniellidae (1)
Xenosauridae (1)
Dipsadidae (7)
Gymnophthalmidae (4)
Crotaphytidae (3)
Lacertidae (78)
Viperidae (35)
Carphodactylidae (6)
Corytophanidae (4)
Pythonidae (9)
Colubridae (41)
Pygopodidae (1)
Psammophiidae (2)
Varanidae (37)
Lamprophiidae (1)
Leptotyphlopidae (1)
Bolyeridae (1)
Cylindrophiidae (1)
Dibamidae (1)
Gerrhosauridae (1)
Blanidae (1)
Shinisauridae (1)
Leiosauridae (1)
0.1
0.5
0.9
1.3
NRMSE
Testudines
Dermochelyidae (1)
Trionychidae (24)
Testudinidae (41)
Emydidae (32)
Platysternidae (1)
Kinosternidae (12)
Podocnemididae (5)
Cheloniidae (5)
Geoemydidae (46)
Carettochelyidae (1)
Pelomedusidae (1)
Chelidae (9)
Chelydridae (3)
0.1
0.3
0.5
0.7
0.9
NRMSE

*Fig S8. Plot showing Normalized Root Mean Square Error of the family block validation for GAM (all groups) and PGLS (squamates and amphibians only). Range intervals represent the NRMSE of GAM and PGLS, while points represent the average value. Numbers near family names represent respective family size. Dashed red line at 0.5 represents the error equal to half of the generation length of the family.*

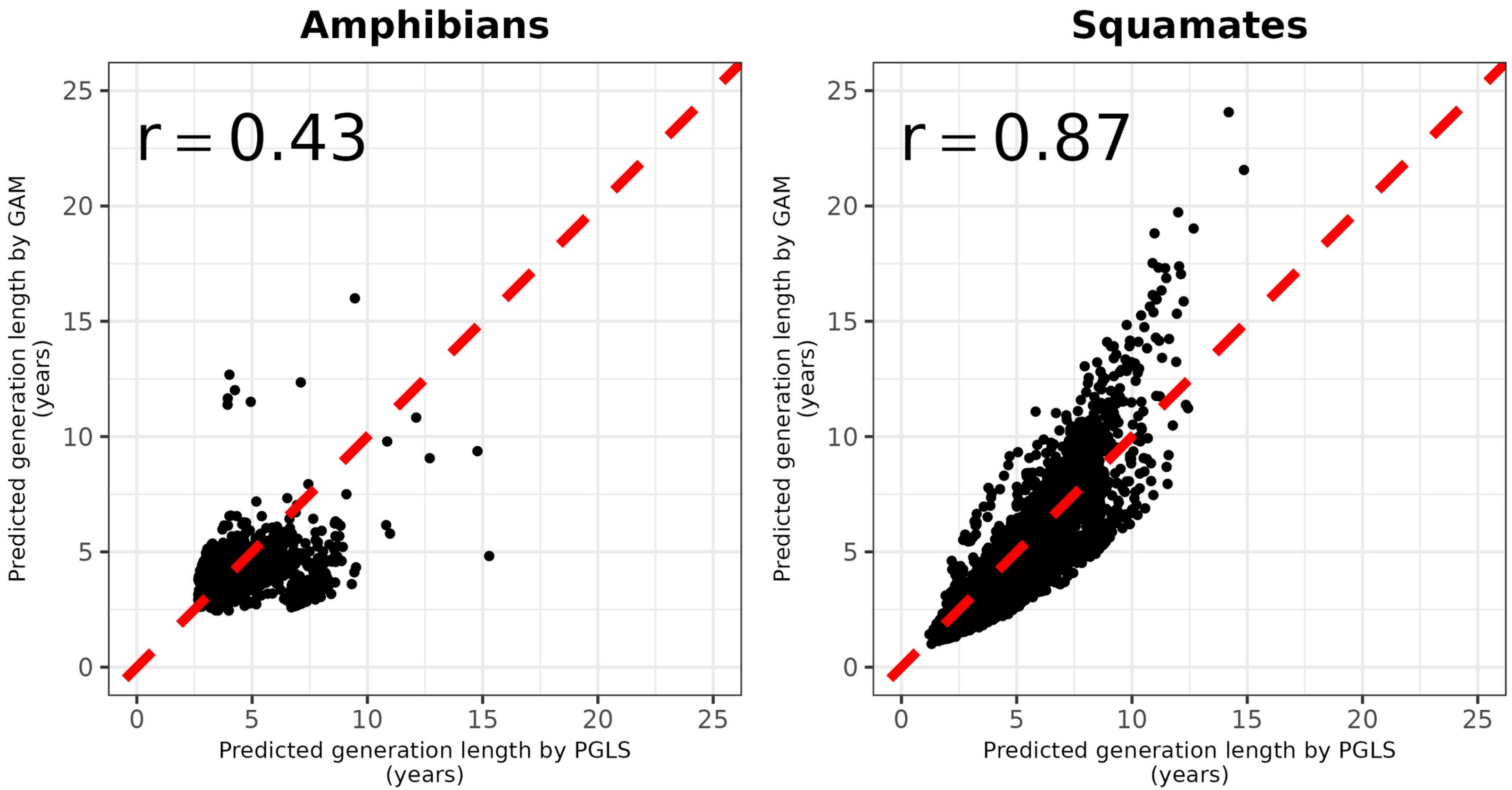


*Fig.S9. Scatter plot showing the relationship between the generation length predicted by PGLS (x axis) and by GAM (y axis) for squamates and amphibians with the Pearson r.*

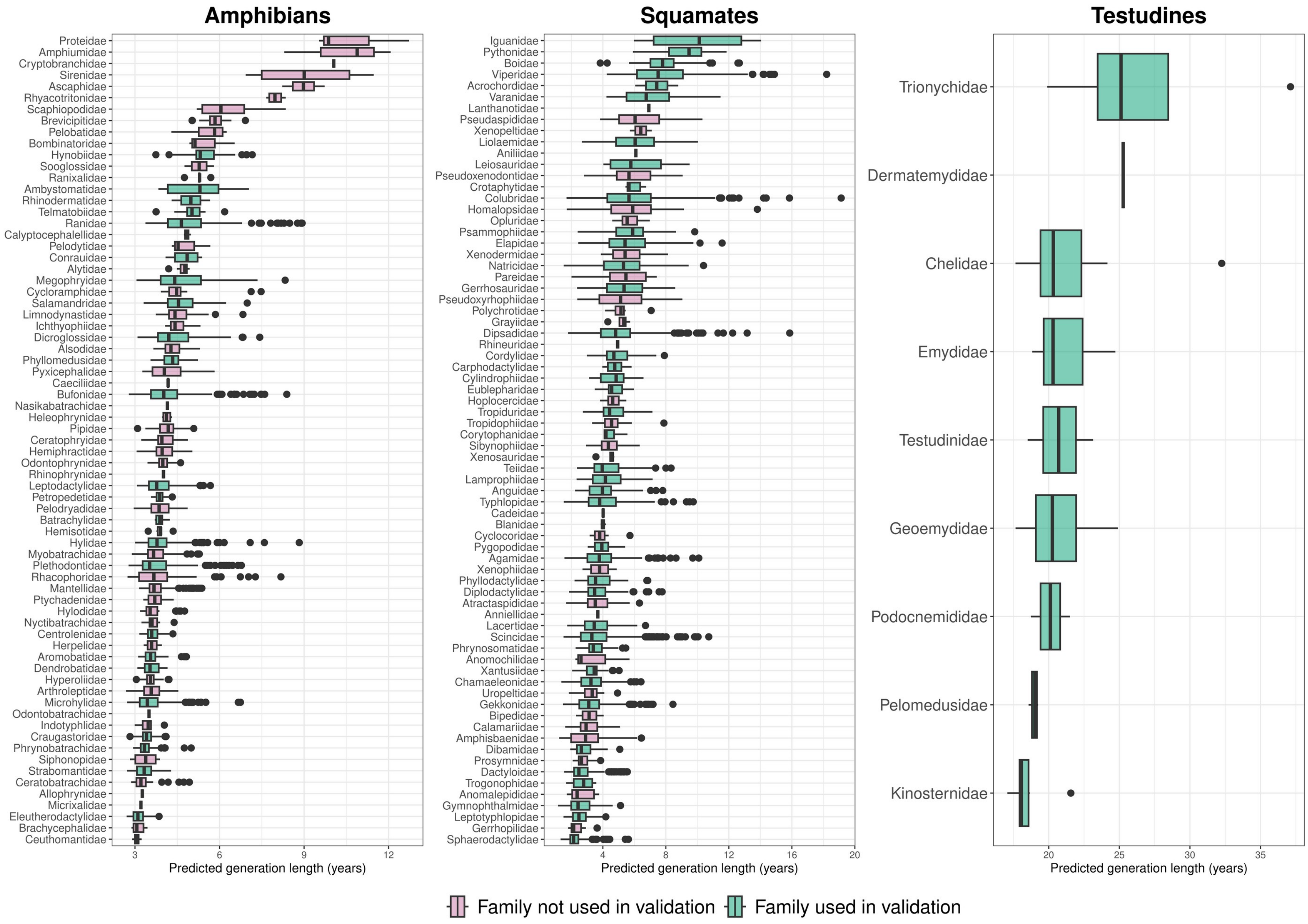

Amphibians
Squamates
Testudines
Predicted generation length (years)
Predicted generation length (years)
Predicted generation length (years)
Trionychidae
Dermatemydidae
Chelidae
Emydidae
Testudinidae
Geoemydidae
Podocnemididae
Pelomedusidae
Kinosternidae
Family not used in validation
Family used in validation

*Fig. S10. Boxplots of the predicted generation length for squamates, testudines and amphibian species. Generation length of squamates and amphibians is the average of the predicted generation length by GAM and PGLS.*

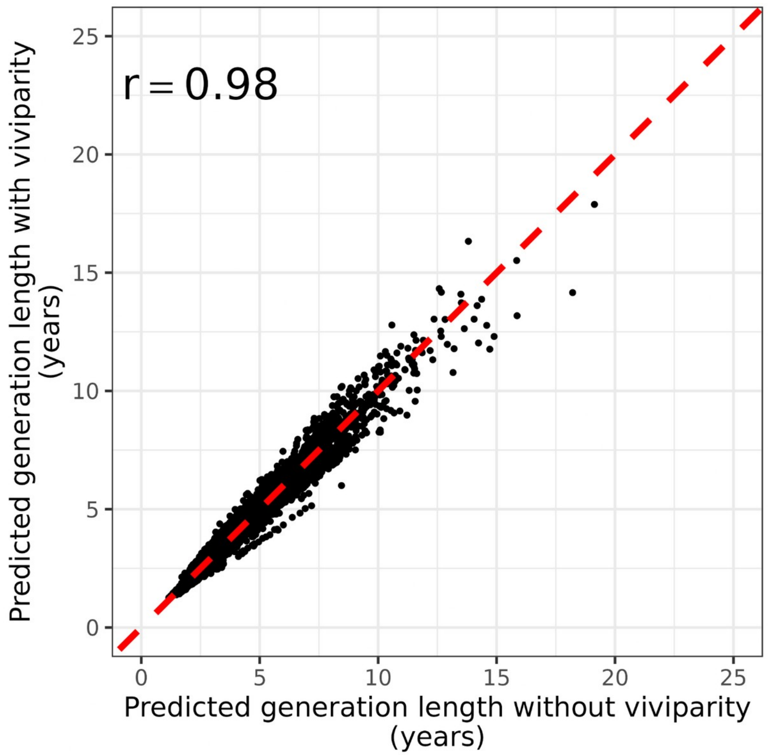


*Fig. S11. Scatter plot showing the relationship between the generation length predicted by without viviparity as a predictor (x axis) and with viviparity as a predictor (y axis) for squamates, with the Pearson r.*

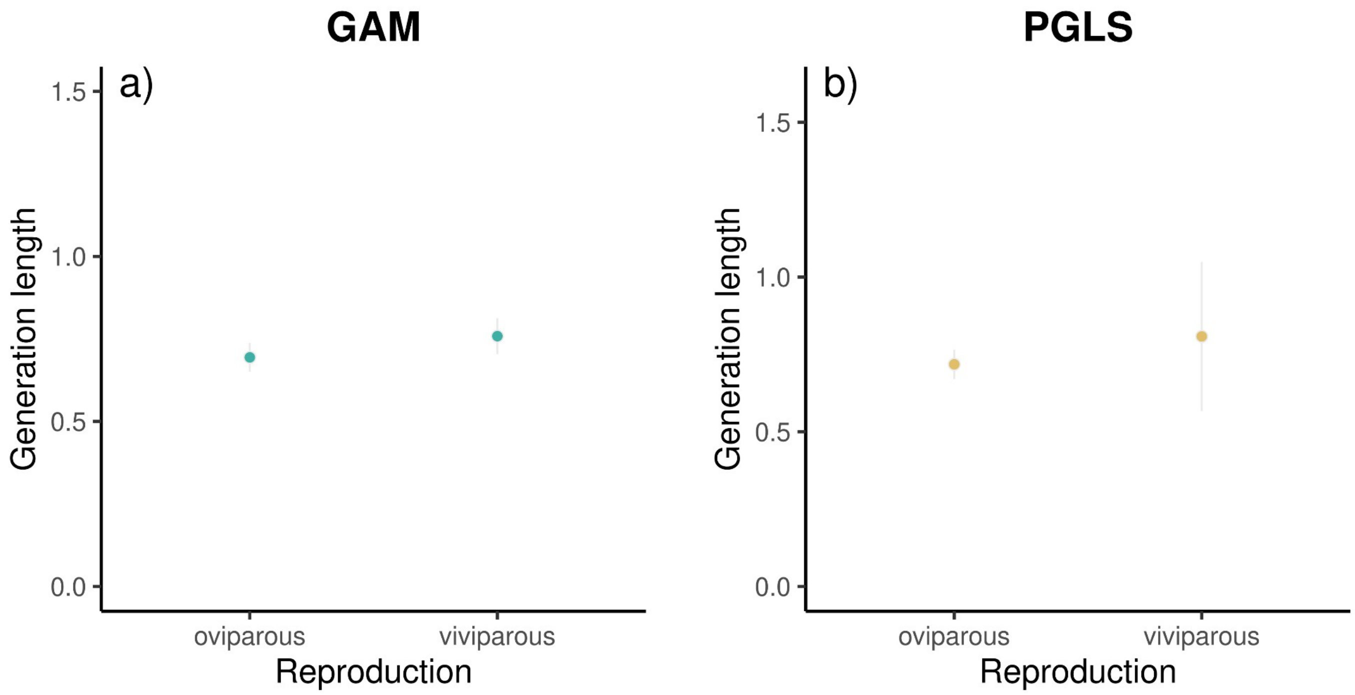


*Fig S12. Partial dependence plots displaying the relationship between generation length and viviparity for squamates in GAM and PGLS. Gray lines represented standard error.*
